# Microtubule polymerization tuned by macromolecular crowdant size and density

**DOI:** 10.1101/2024.02.02.578534

**Authors:** Jashaswi Basu, Aman Soni, Chaitanya A. Athale

## Abstract

Microtubule (MT) polymerization is regulated by biochemical as well as physical factors such as macromolecular crowding. Crowding agents or crowdants affect MT elongation rates differently depending on crowdant size due to opposing effects on polymerization: microviscosity reduces polymer elongation, while volume exclusion increases reaction rates by local concentration. In order to address how crowdant size and concentration collectively affect MT populations, we combine *in vitro* MT polymerization experiments with kinetic Monte Carlo simulations. Our experiments in bulk with nucleators validate decreasing MT elongation rates with increasing concentrations of small molecular weight crowdants in bulk assays and a corresponding increase for large crowdants. Kinetic Monte Carlo simulations can explain the result with packing fractions dependence of small as compared to large crowdants increasing microviscosity more dramatically. In contrast MT bulk polymerization rates in absence of nucleators increased with crowdant concentration, irrespective of their size, with a corresponding decrease in the critical concentration. Microscopy of filament growth dynamics demonstrates that small crowdants result in shorter filaments in a concentration dependent manner, consistent with their role in reducing elongation rates, but this decrease is compensated by increased number of filaments. Large crowdants increase the filament numbers while elongation is slightly decreased. Our results provide evidence for MT nucleation being rate-limited and elongation diffusion limited, resulting in differences in the effect of crowdant sizes on nucleation and elongation. These results are of general relevance to understand physical effects of crowding on collective cytoskeletal polymerization dynamics.

## Introduction

Microtubules (MTs) are intracellular cytoskeletal filaments whose polymerization from *α* and *β* tubulin monomers are vital for cell growth and physiology. The polymerization dynamics of MTs are described by nucleation dependent kinetics (NDP) model^1,2^, with monomers aggregating above a critical concentration (c*), to form stable nuclei (MT nucleation). MT elongation occurs by the addition of tubulin subunits by GTP hydrolysis ^3^. At the same time at a single filament level, MTs show ‘dynamic instability’ defined as the transition between growing and shrinking states^4^. This has been modeled by a two state four parameter model capturing the transition frequencies of growing to shrinking -catastrophe - and shrinking to growing -rescue -together with the growth and shrinkage speeds^5^. This model however assumes filaments to be independent of each other. Most experimental studies of MT polymerization dynamics address either dynamics in bulk scattering assays^6^, or at a single filament level using total internal reflection (TIRF) microscopy^7^ or interference reflection microscopy (IRM)^8^. However, polymerizing tubulin involves a population of MTs whose numbers and sizes are fluctuating influenced by monomers diffusion and interaction with the emergence of multiple filaments dynamics. Such collective effects are more closely related to the dynamics in cells, but are less well understood.

One such collective effect that has been studied widely in the context of transport and enzyme kinetics is macromolecular crowding (MMC), that results from a crowded and spatially structured cytoplasmic environment seen *in vivo*. MMC is described as the reduction in available volume for reactants due to the concentration of macromolecules occupying a significant fraction of the cell^9^. The effects are varied ranging from excluded volume, depletion effects to viscosity changes^10^, that in turn can affect cell physiology^11^. Crowding agent size and concentration have been previously reported to influence reaction rates in both experiment and theory as seen in case of the significant decrease in association rates of the *β*-Lactamase (TEM-1) and *β*-Lactamase inhibitor protein (BLIP) system with concentration in presence of small-sized crowdants but less so for large crowdants^12,13^. Amyloid fibre association rates in contrast were enhanced by small molecular weight crowding agents in a concentration dependent manner^14^. Simulations have been used to disambiguate these differential outcomes based on the nature of the association reaction, with the rates of diffusion limited reactions decreased due to crowdants as a result of reduced mobility - viscogenic effect - while they were increased if the process was rate limited - excluded volume effect - assuming crowdant and interactor sizes are the same^15^. Simulations attempting to mimic cytoplasmic conditions with a distribution of crowdant molecule sizes demonstrated that low packing fractions of crowdants increased association rates through excluded volume while higher packing fractions reduced the rates through viscogenic effects ^16^. In addi-tion, qualitative differences in spatial structure imposed by crowdants, localized microdomains or corrals affected quantitative association dynamics of receptors at steady state in simulations^17^. Models of cytoskeletal filament polymerization in the context of the combined effect of crowdant size and concentration however remain less well studied.

In experimental work with actin cytoskeletal filament elongation rates (in presence of ‘seeds’) it was found that, low molecular weight (LMW) crowdants slow them down, while high molecular weight (HMW) crowdants increased rates^18^. In contrast, the rate of *de novo* bulk polymerization of actin in the absence of nucleating ‘seeds’ was unaffected by LMW crowdants, while HMW crowdants increased rates^19^. A comparable study on single filament MT polymerization dynamics *in vitro* has demonstrated that LMW crowdants resulted in a decrease of elongation rates, while HMW crowdants increased them, in a concentration dependent manner^20^. An *in vivo* study demonstrated osmotic stress induced increases in cytoplasmic concentration resulted in a reduction in MT growth rates in *Schizosaccharomyces pombe* cells that were correlated with the increased viscosity, comparable to *in vitro* effects of glycerol as a crowdant on MT elongation^21^. Therefore MT and in general cytoskeletal polymerization dynamics are affected by crowdants in a size-dependent manner. Since crowdants can act by multiple mechanisms, the sometimes contradictory data is in need of a consistent explanation both in terms of experiment and simulations.

Computational modeling of physical mechanisms of regulation of MTs have successfully predicted the emergence of bounded and unbounded phases of growth as a mechanism of MT regulation^5^ as well as the role of the GTP-cap in determining catastrophe frequencies observed experimentally in dilution experiments^22^. Kinetic models of the nucleation-dependent polymerization type with four-step nucleation were fit to bulk kinetic polymerization data resulting in an estimate of the nucleating ‘seed’ size^23^. Kinetic Monte Carlo simulations of the effect of monomer diffusion rates on MT growth demonstrated oscillations and length distributions in an aster influenced by monomer diffusion limited growth as compared to fast diffusing monomers^24^. Brownian Dynamics simulations have shown increase in association rate arising from excluded volume interactions^25^ while experiments show MT elongation to be diffusion limited^20^. A computational model that integrates kinetics of growth of MTs and their effect on monomer diffusion and MT elongation could help better interpret experimentally observed effects of crowdant size and concentration.

Here, we have examined the physical effects of crowdant size and concentration on MT polymerization dynamics using a combination of *in vitro* experiments, microscopy and stochastic simulations. We show that elongation rates of MT filaments from nucleating ‘seeds’ are decreased by low molecular weight (LMW) crow-dants, and increased by high molecular weight (HMW) crowdants in turbidimetry. We compare these experimental results to a lattice-based kinetic Monte Carlo simulation of monomer diffusion and filament elongation from ‘seeds’ in presence of crowdants of increasing size. Simulations show reduced elongation rates in presence of increasing crowdant concentrations, with crowdants smaller in size to monomers resulting in more dramatic decrease as compared to larger. Polymerization of MTs in the absence of ‘seeds’ that involves nucleation and elongation, i.e. *de novo* polymerization kinetic rates in contrast appear to increase with crowdant concentration in a size-independent manner. By measuring the effect of crowdants on the critical concentration from bulk turbidimetry and filament lengths and numbers from microscopy, we examine how crowdant concentration, viscosity and size affect MT elongation and nucleation. These findings demonstrate that the effect of crowding varies from viscogenic to excluded volume and in the absence of nucleators regulates MT polymerization by physical mechanisms.

## Results

### Effect of crowdant size and concentration on bulk microtubule elongation dynamics

Increased osmotic stress *in vivo* and glycerol concentration *in vitro* were seen in previous work to reduce single MT filament elongation rates, attributed to increase in viscosity^21^. At the same time multiple crowdants were shown to uniformly increasing viscosity with concentration, but had contrary effects on single filament MT elongation rates depending on crowdant size - decreasing with glycerol and ethylene glycol, while increasing with PEG, BSA and dextran^20^. We proceeded to examine if collective polymerization, i.e. in bulk turbidimetry would also show the same crowdant size-dependent effect. Tubulin polymerization in presence of GMPCPP stabilized MT ‘seeds’ assembled from 30 *µ*M tubulin were combined with low concentrations of tubulin monomers (8 *µ*M), below the critical concentration in a volumetric 5:3 ratio (details in the Materials and Methods). The mixture was combined with a crowding agent-either glycerol, PEG 4000 or Ficoll 400 and absorbance at 340 nm used to measure polymerization (Figure 1A).

**Figure 1:**
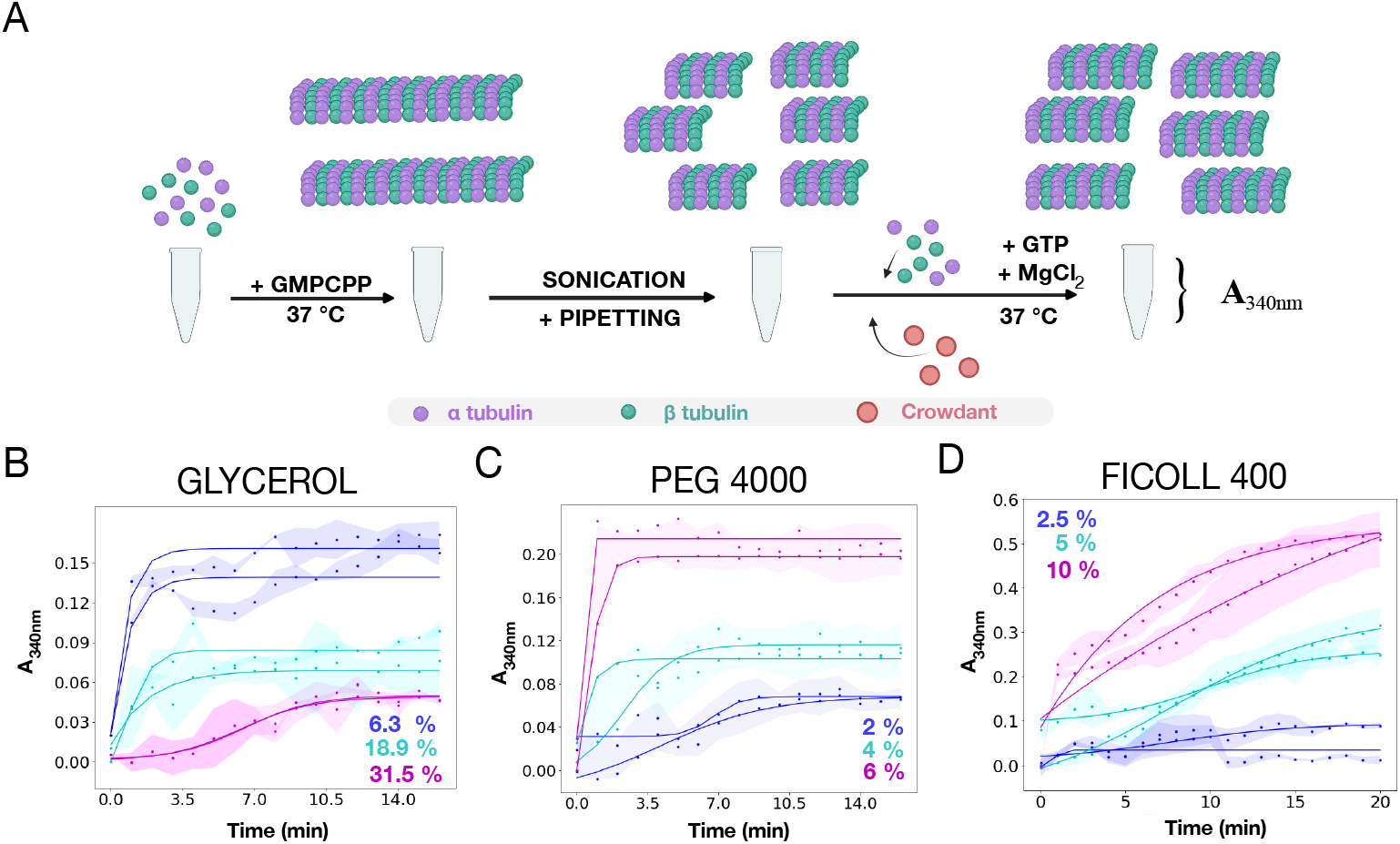
Effect of crowdant molecular weight and concentration on mi-crotubule (MT) elongation kinetics. (A) The schematic depicts the steps involved in MT-seed nucleated polymerization kinetics with sheared, GMPCPP-stabilized MTs serving as seeds for GTP-dependent tubulin polymerization (elon-gation) in presence of crowdants measured as absorbance kinetics (further details are in the Materials and Methods section). (B-D) The plots represent the ab-sorbance at 340 nm (A_340_*_nm_*) measured as a function of time in minutes, of a system containing GMPCPP-stabilized MT seeds with the addition of 8 *µ*M goat brain tubulin and 1 mM GTP being constant in presence of three macromolecular crowdants (MMCs): (B) glycerol (92 Da), (C) PEG 4000 (4×10^3^ Da) or (D) Ficoll 400 (4×10^5^ Da). MMC concentration is reported as %w/v (colors). Colored circles: mean absorbance, shaded region: standard deviation (n = 9), solid lines: fit line based on the nucleation dependent polymerization (NDP) model (Equation 1).

Crowdant concentrations affect polymerization both in terms of steady state absorbance at saturation, reflecting the polymer mass fraction, and rate of increase in absorbance, a measure of polymerization rate. While glycerol decreases both polymerization rate and steady state polymer mass (Figure 1B), PEG 4000 (Figure 1C) and Ficoll 400 (Figure 1D), increased both polymer mass and rate of polymerization in a concentration dependent manner.

We quantify the observations by fitting a nucleation dependent polymerization (NDP) kinetic model (Equation 1) to the absorbance kinetic data, that has been previously used to characterize MT polymerization kinetics by turbidimetry^23,26^. The model is used to estimate three parameters: the normalized steady state absorbance [A*_max_* - A_0_] (where A*_max_*: steady state absorbance, A_0_: initial value), the polymerization rate *r* and half maximal time of polymerization t_1_*_/_*_2_. Consistent with the qualitative observations, of a size dependence, we observe that increasing glycerol concentration results in decreasing steady state absorbance (Figure 2A(i)) and growth rates (Figure 2A(ii)) while increasing the t_1_*_/_*_2_ (Figure 2A(iii)). In contrast, PEG-4000 and Ficoll 400 both result in increasing relative steady state absorbances (Figure 2B,C(i)) and polymerization rates (Figure 2B,C(ii)), while decreasing the half-maximal time (Figure 2B,C(iii)).

**Figure 2:**
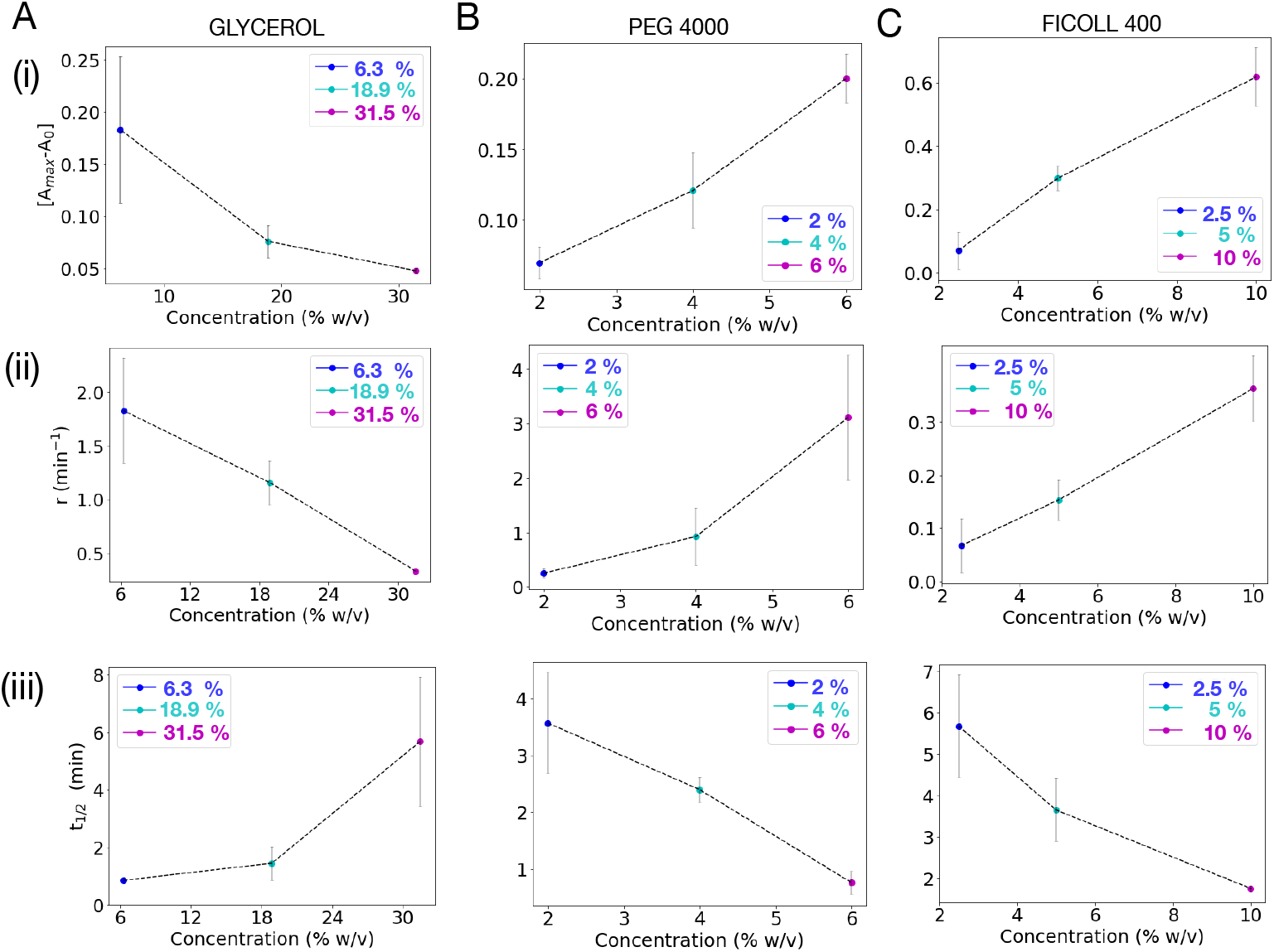
Crowdant molecular weight determines concentration depen-dence of MT elongation rates. (i-iii) The effect of crowdant concentration (x-axis) on the following fit parameters of the seed-nucleated MT elongation are plotted: (i) [*A_max_ −A*_0_]: normalized steady state absorbance, (ii) r: polymerization rate (min*^−^*^1^) and (iii) *t*_1_*_/_*_2_: time for the kinetics to reach half-maximal absorbance (min). (A-C) Increasing concentrations (%w/v) of the following crowdants were used: (A) Glycerol, (B) PEG 4000 and (C) Ficoll 400. Error bars: Standard deviation from 2 biological replicates with three technical replicates each (n = 6).

This demonstrates MT elongation rate and steady state mass are both reduced by increasing glycerol concentration, but increased by PEG-4000 and Ficoll 400. We find the effect of crowdant size on collective polymerization is consistent with previously reported effects using single filament dynamics^20^. Previously, the reduced elongation rates had been attributed to viscosity *in vivo*^21^. However, LMW and HMW crowdants both increase bulk viscosity as we see in our measurements for glycerol and PEG 4000 (Figure S1) and reported elsewhere^19^. We therefore proceeded to examine what mechanism could determine the difference in the effect of crowdants on polymerization in presence of seeds, using a computational model.

### Kinetic Monte Carlo model predicts role of microviscosity in crowdant size dependence on MT elongation

A computational model of was developed that combines (a) *diffusion model* of monomers and crowdants, whose size is varied with (b) a *polymerization model* of monomers stochastically assembling to form linear filaments, a detailed description of which can be found in the Materials and Methods section. As a fist step, simulations tested the effect of crowdants of increasing size and packing fraction on the diffusion of monomers. In the ‘toy model’ the molecular weight of crowdants (LMW and HMW) was mapped to size in terms of lattice area, while monomers were maintained at a fixed size. Therefore monomers were modeled to occupy-ing a square of side length 3 lattice units i.e. *L_monomer_* = 3, while crowdants occupied squares with side-lengths *L_crowdant_* of 1, 3, 5, 7 or 9 (Table S1). This approach allowed us to examine the effects of both increasing crowdant size and packing fraction, i.e. relative area occupied by the crowdant. Monomer density was maintained constant in all calculations. Diffusive trajectories of monomers over 20 seconds can be seen qualitatively to be confined in presence of small crow-dants (LMW) as compared to the wider spread of movement in presence of large (HMW) crowdants (Figure S2). To quantitatively assess these observations, we calculated the diffusion coefficients of the tracks from MSD plots (Figure 3A, as described in the Materials and Methods section). We find that the effective diffusion coefficient (*D_eff_*) decreases for all crowdant types, but is more pronounced for small (LMW) crowdants as compared to large (HMW) crowdants (Figure 3D). The exponent of anomalous diffusion, *α*, decreases with packing fraction for all crowdants (Figure S3A). This suggests that in general the mobility shows a tendency to become increasingly sub-diffusive, a trend also reflected in the changing slope of the MSD curves for Δ*t* greater than *∼*1 second for crowdants larger than 3 (Figure 3A). Polymerization kinetics were simulated in presence of pre-existing nucleators or ‘seeds’, consisting of multiple monomers denoted by *N_s_* and freely diffusible monomers (Table S1). The effect of increasing crowdant packing fraction (*ϕ_C_*) was to reduce the slope and maximal value in case of small crowdants (*L_crowdant_* =1 and 3) (Figure 3B, *left*), while large crowdants (*L_crowdant_*= 5, 7, 9) had a less prominent effect on polymerization (Figure 3B, *right*). These results suggest that small sized crowdants affect diffusion and polymerization through the increase in effective microviscosity, while for the same packing fraction large sized crowdants over short times appear to have little or no effect. The effect on microviscosity is schematically depicted to imply that at short time scales large crowdants at the same packing fraction result in regions devoid of crowdants and allow for free diffusion of monomers, while the same packing fraction of small crow-dants, monomers interact more frequently with crowdant molecules thus reducing their mobility (Figure 3C), as demonstrated by the effect of crowdant size and packing fraction on the effective diffusion coefficient of monomers (Figure 3D). The steady state polymer fraction shows a drastic decrease with crowdant packing fractions for small (*L_crowdant_*=1), but little or no change for all other crowdants from *L_crowdant_*=3 to 9 (Figure 3E). Their effect on MT lengths is consistent with the effect on diffusion coefficients (Figure S3B). These results are consistent both with our experimental kinetics of MT elongation described in the previous section in terms of the effect of small crowdants (1×1) but do not shown an increase in polymer fraction or mean length for equal sized or larger crowdants (3×3 and larger), that is observed in experiment. This could possibly result from the weaker effect of volume exclusion as expected in our observations and those of others, and suggests the need for further exploration of the model dynamics and their parameter dependence.

**Figure 3:**
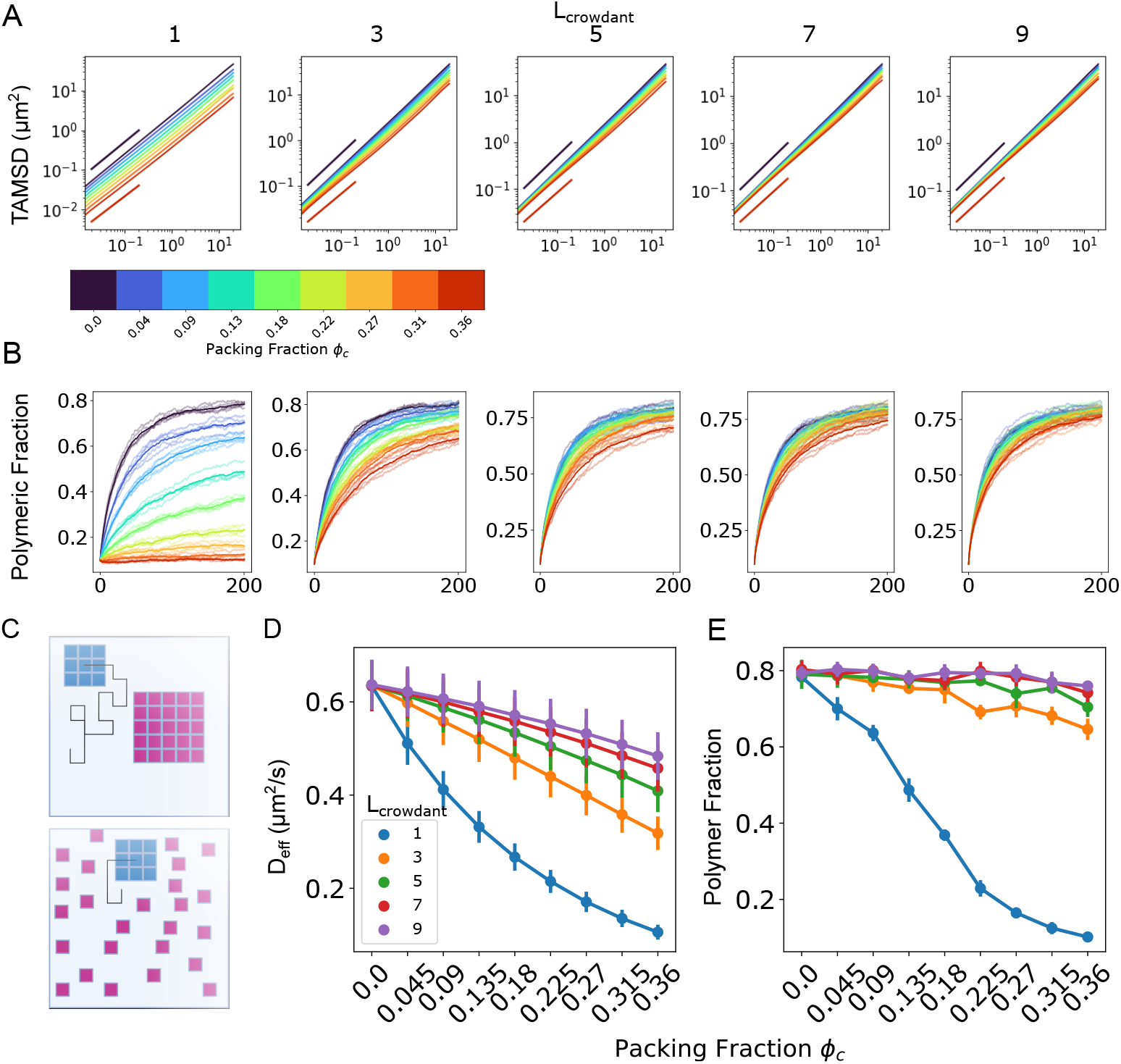
Simulating the effect of crowdant size and density on monomer diffusion and filament polymerization. (A) Simulated XY trajectories of monomers (size 3×3) were used to calculate the time averaged mean square dis-placement (TAMSD) and plotted on a log-log plot as a function of time interval (x-axis). Crowdant sizes, *L_crowdant_*, were varied (left to right): 1, 3, 5, 7 and 9, while packing fractions ranged from 0 to 0.36 (color bar). The solid short lines denote regions used for fitting. (B) Simulations kinetics of polymer growth in pres-ence of pre-existing seeds (1:10 seeds:monomers) shows the total polymer fraction (y-axis) increase with time (x-axis). Color bar: area packing fraction. (C) The schematic represents an explanation for the differential effect of small and large sized crowdants on the monomer mobility, for the same area packing fraction. (D,E) The crowdant packing fraction *ϕ_c_* dependence (x-axis) of (D) the effective diffusion coefficient (y-axis) from purely diffusive simulations obtained from fitting the anomalous diffusion model and (E) the fractional polymer mass at the end of simulations indicated in (B) are plotted as a function of packing fraction. Colors: crowdant size.

MTs in the cellular context are likely to also spontaneously nucleate based on the reported *in vivo* tubulin cellular concentration of *∼*2mg/ml^27,28^, since it exceeds the critical concentration. We thus proceeded to examine how the combined process of nucleation and elongation are affected by crowdant size and concentration.

### Crowdant concentration increases *de novo* polymerization independent of crowdant size

Microtubule polymerization is nucleation-dependent and involves two steps, a rate limited MT nucleation^3^ and elongation, together referred to as nucleation dependent polymerization, NDP^23,26,29,30^. We proceeded to test the effect of crowdants on both nucleation and elongation by polymerizing 30 *µ*M tubulin in the presence of 1 mM GTP and increasing concentrations of crowdants without the addition of seeds, as schematically depicted (Figure 4A). This has been previously referred to as *de novo* polymerization^31^. The absorbance kinetics for both LMW (glycerol) as well as HMW (PEG 4000 and Ficoll 400) all appear to show three phases in their kinetics i.e lag-, growth- and saturation-phase, irrespective of crowdant size (Figure 4B-D). We observe a common effect of decreased lag periods and increased steady state absorbance with increasing crowdant concentration, in glycerol (Figure 4B), PEG4000 (Figure 4C) and Ficoll 400 (Figure 4D). In addition, another LMW crowdant ethylene glycol (MW: 62 Da) (Figure S4A) as well as multiple HMW crowdants such as PEG 20,000 (MW: 20 kDa) (Figure S4B), bovine serum albumin, BSA (MW: 66 kDa) (Figure S4C) and methylcellulose (14 kDa) (Figure S4D) all resulted in increased rate and steady state absorbance.

**Figure 4:**
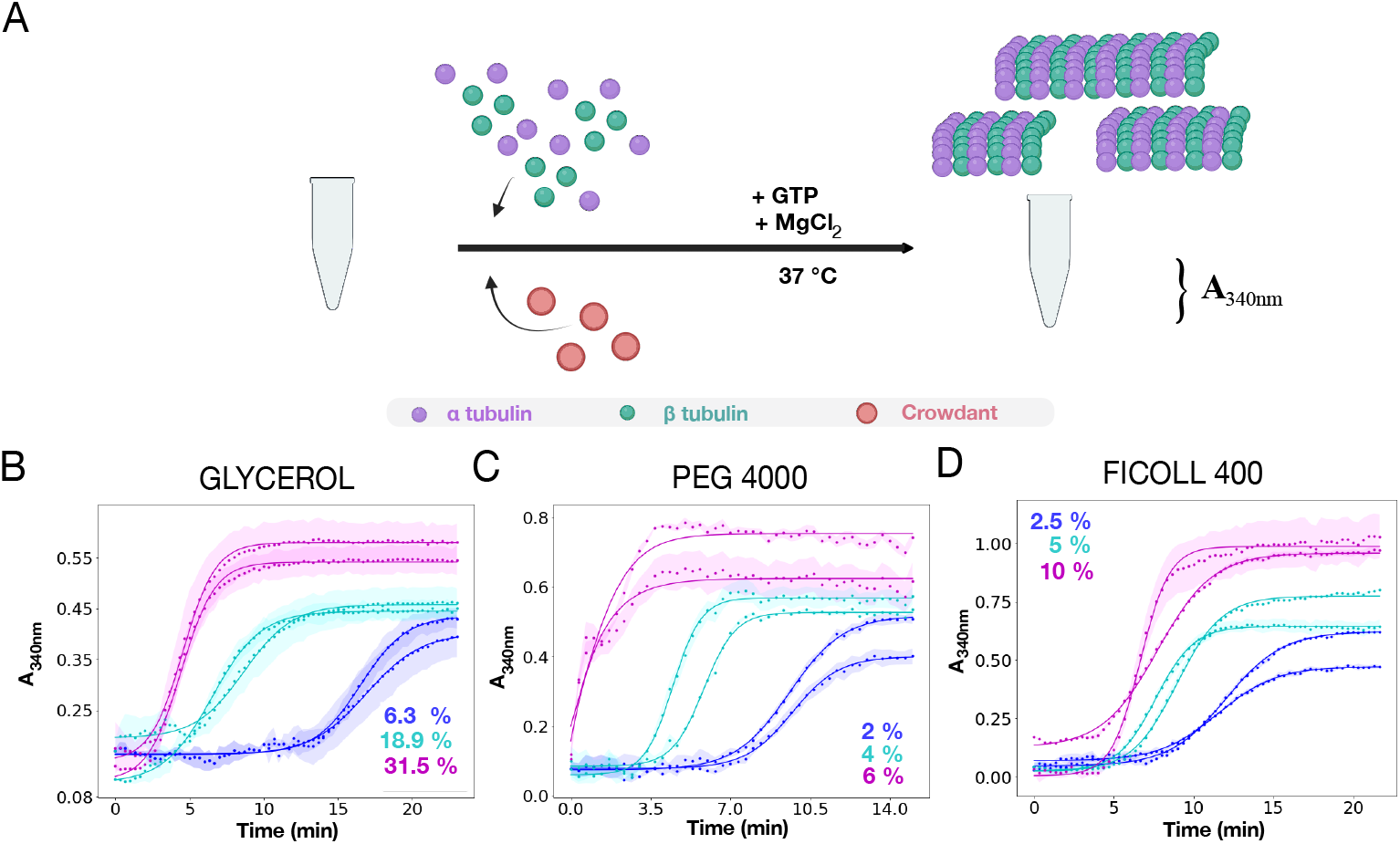
Effect of crowdant molecular weight and concentration on the kinetics of *d* e novo MT polymerization. (A) The schematic represents how polymerization of *α*, *β* tubulin monomers (green and purple circles) was followed in presence of crowdants (red circles) by absorbance kinetics. The experiments were conducted without the addition of nucleating seeds. (B-D) The absorbance at 340 nm (A_340_*_nm_*) is plotted as a function of time in minutes, as a measure of the polymerization of 30 *µ*M goat brain tubulin with 1 mM GTP, with increasing concentrations (%w/v) of three different crowdants: (B) glycerol (MW 92 Da), (C) PEG 4000 (4×10^3^ Da) or (D) Ficoll 400 (4×10^5^ Da). The plot depicts mean (n = 6) absorbance (colored circles) with the standard deviation of each biological replicates (shaded region). Solid lines: fit line based on the NDP model (Equation 1). Colors: crowdant concentrations.

In order to examine the quantitative relation between crowdant sizes and concentrations in terms of their effect on polymerization mass and rates, we used the same NDP model (Equation 1) to fit our data, as before with fit parameters: steady state absorbance [A*_max_*-A_0_], polymerization rate (r) and time to reach half-maximal absorbance (t_1_*_/_*_2_). Consistent to our qualitative observation, increasing concentrations of all the three crowdants - glycerol, PEG 4000 and Ficoll 400, resulted in increasing [A*_max_*-A_0_] (Figure 5A-C (i)) and r (Figure 5A-C (ii)) and a corresponding decrease in t_1_*_/_*_2_ (Figure 5A-C (iii)) respectively. This demonstrates that crowdants irrespective of their size (LMW as well as HMW) result in a concentration dependent increase in polymerization kinetics. This is the observed when both MT nucleation and elongation are simultaneously acting to increase polymer mass, i.e. *de novo* polymerization.

**Figure 5:**
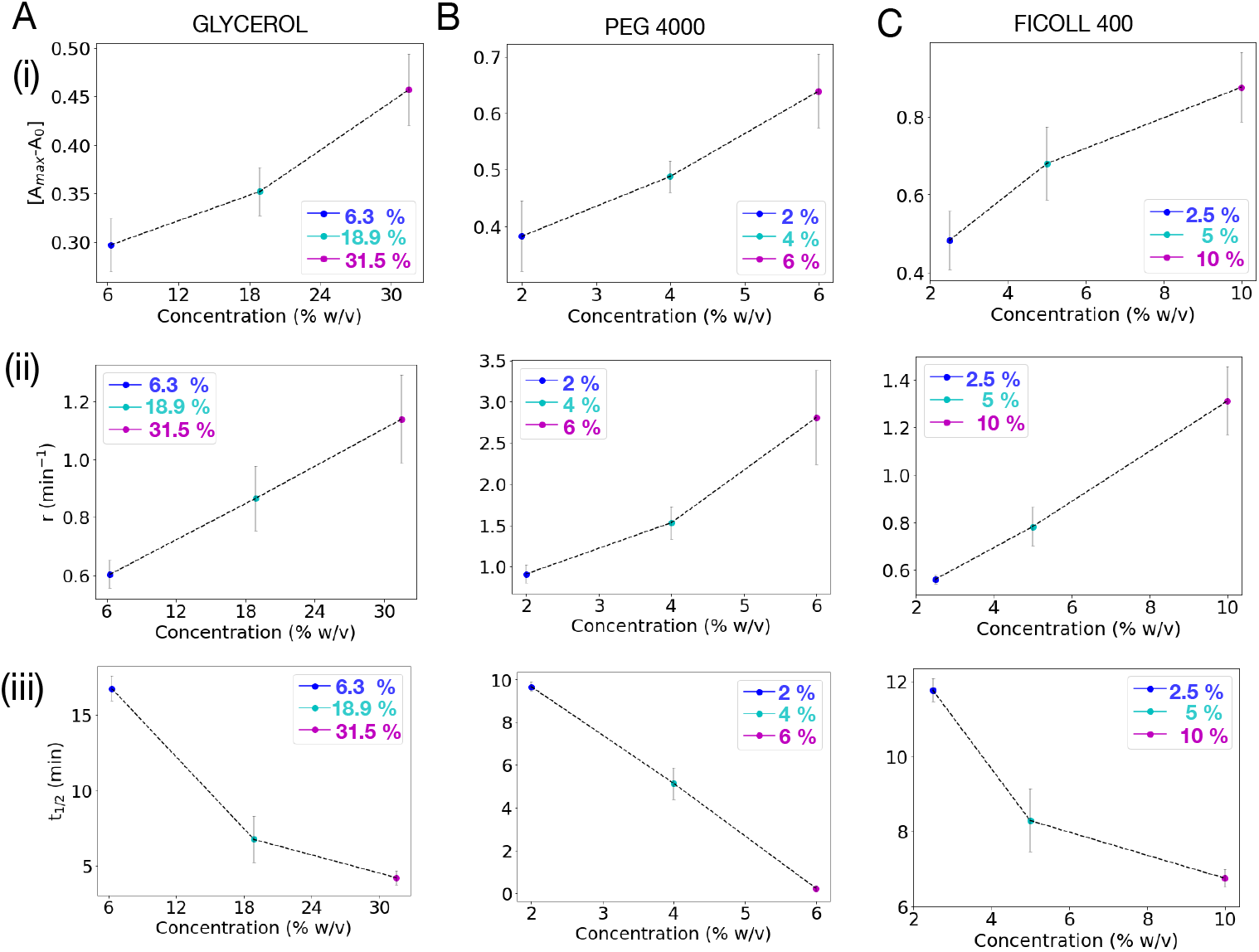
*De novo* MT polymerization rates dependence on crowdant con-centration is independent of molecular weight. The effect of crowdants obtained from fitting kinetic data to NDP model (Equation 1) are quantified in terms of (y-axis): (i) [A*_max_*-A_0_]: the normalized steady state absorbance, (ii) r: polymerization rate (min*^−^*^1^) and (iii) t_1_*_/_*_2_: time corresponding to half-maximal absorbance (min). Data is plotted as a function of increasing concentration %w/v (x-axis) of three different crowdants: (A) glycerol (B) PEG 4000 and (C) Ficoll 400. Circles: Averages, error bars: s.d. from 2 biological replicates with three technical replicates each (n = 6).

The differential effect of crowdant size on bulk MT elongation and *de novo* MT polymerization, suggests crowding affects nucleation differently than MT elongation. To understand this, we measured the critical concentration of tubulin in presence of crowdants.

### Crowdant size independent decrease of MT critical concentration of nucleation, independent of crowdant size

Our observations suggest that the effect of crowdant size differs between MT elongation rates and *de novo* polymerization. In order to understand the mechanism of differences in the effect of crowding on nucleation as compared to elongation, we proceeded to measure the critical concentration (c*) of tubulin as a readout of MT nucleation^1,26,32^. The kinetics of *de novo* polymerization were measured in the presence of increasing concentrations of LMW (glycerol) and HMW (PEG 4000) crowdants, for a fixed range of tubulin monomers (10 to 30 *µ*M). The normalized steady state absorbance ([A*_max_*-A_0_]) were measured as a representative of polymer mass and plotted as a function of monomer concentration (Figure 6A, B). Based on the definition that critical concentration, c*, is that monomer concentration, below which the steady state polymer fraction is zero, we fit a straight line to the data points and estimated c* from the x-intercept. We find c* reduces with crow-dant concentration for both glycerol (Figure 6C) and PEG 4000 (Figure 6D). We find the critical concentration of goat tubulin is reduced by a factor of 2 over the range of concentrations tested of both glycerol and PEG (Table S2), compared to 5 *µ*M reported without the addition of crowdants^26^. This suggests MT nucleation is enhanced by increasing crowdant concentration, independent of its size.

**Figure 6:**
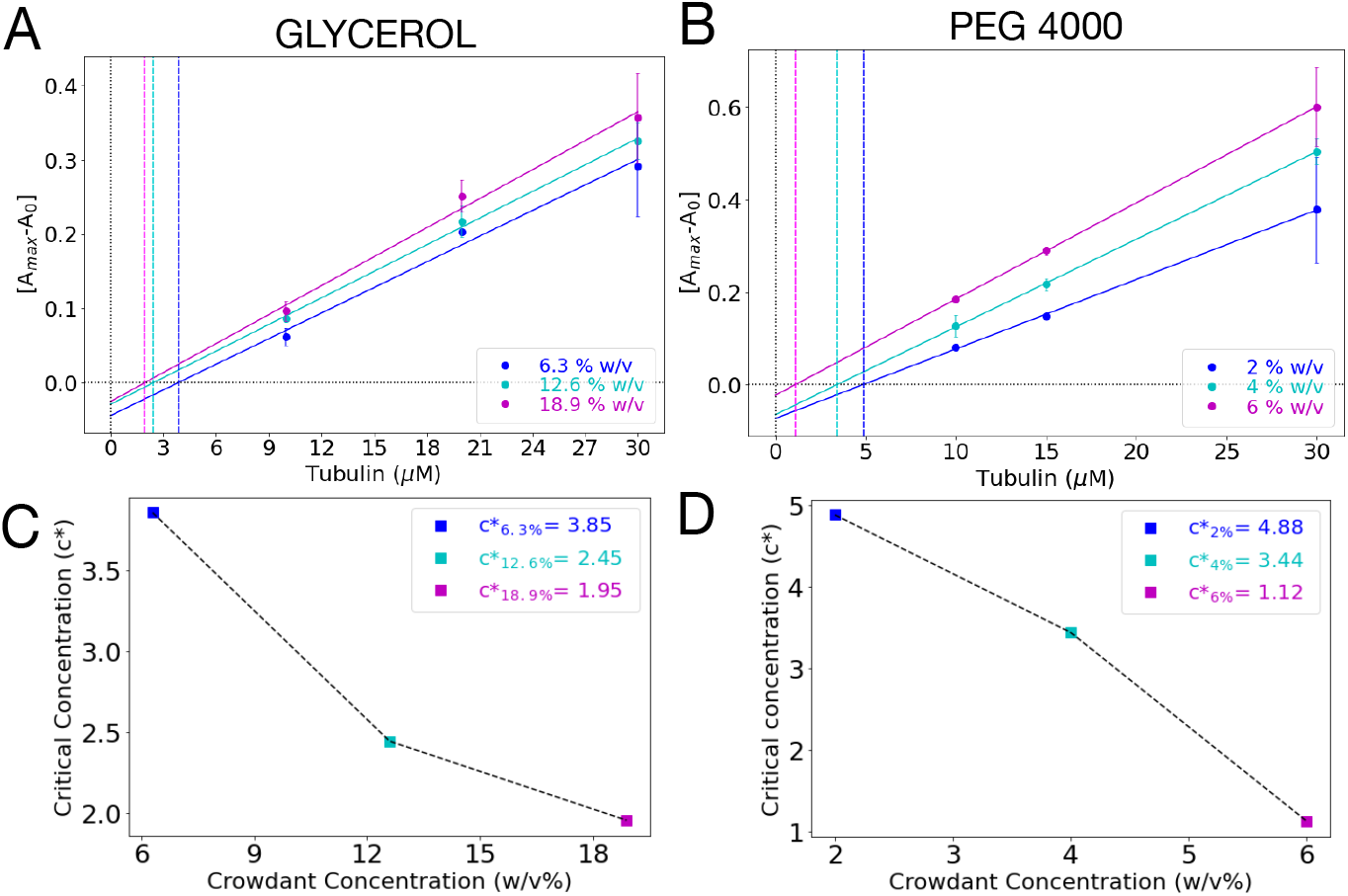
Effect of glycerol and PEG 4000 on critical concentration of tubulin polymerization. (A, B) The normalized steady state absorbance *A_n_* = [*A_max_ − A*_0_], representing the polymer mass (y-axis) for increasing tubulin concentrations (x-axis) is plotted with increasing (A) glycerol and (B) PEG 4000. Circles with errorbars: mean*±*s.d. (n = 6), colors: crowdant concentration. The values were then fit by a straight line model (Equation 2). The x-intercept of the line is the critical concentration c* (dashed lines). (C, D) The critical concentration (y-axis) is plotted against different concentrations (% w/v) (x-axis) of the crowdants (C) glycerol and (D) PEG 4000.

Thus, from our *de novo* MT polymerization, we conclude MT nucleation appears to be independent of crowdant size, depending solely on the concentration of crowdants. However, our current estimations of crowdant size dependence on both nucleation and elongation are based on indirect turbidimetric assay, i.e. they measure total polymer mass. In order to reconcile our findings about elongation being differentially affected by crowdants based on their size, but size independent nucleation, we proceeded to examine *de novo* MT polymerization at a single filament level.

### Increasing MT numbers compensates for decrease in MT length for LMW crowdants

Single filament dynamics are typically are measured one filament at a time and have provided deep insights into regulation of MT polymerization. Here, we wanted to examine the role of possible collective effects, in order to better understand our bulk polymerization kinetics measurements. Tubulin samples with GTP and crowdants were added to double-backed tape chambers and sealed between a slide and coverslip (Figure S5A). We observed emergence of MT filaments from free monomers (nucleation) and the growth of those filaments (elongation) with time in presence of increasing concentrations of either glycerol or PEG 4000 in time lapse label free interference reflection microscopy, IRM (Figure S5B, S6), as described in detail in the Materials and Methods section Both glycerol and PEG 4000 concentration increases result in increased nucleation as seen in the increased filament density after 12 minutes of incubation (Figure 7A,D). We sampled filament lengths after 12 minutes since that was the approximate time taken for sat-uration of *de novo* bulk polymerization kinetics (Figure 4B-D). Filament lengths appear to decrease with increasing concentration of glycerol but not significantly for PEG 4000 (Figure 7B,E). While absolute concentration ranges of glycerol and PEG 4000 differ, the volume packing fractions calculated from their hydrodynamic radii (Table S3) are of similar range (Figure S7). The corresponding mean MT lengths and numbers confirm the qualitative impression that while both increasing concentrations of glycerol and PEG 4000 result in increasing MT numbers - nucleation, glycerol also results in correspondingly shorter filaments - elongation (Figure 7C,F). The decrease in MT lengths for glycerol across concentrations was found to be statistically significant. These results are consistent with both the observed crowdant concentration dependent, but size independent lowering of the critical concentrations i.e. LMW and HMW crowdants both have the same qualitative effect (Figure 6). At the same time, reduction in MT length with increasing glycerol is consistent with the observed slower elongation rate with increasing concentration of LWM crowdants in seed-based polymerization (Figure 2(ii)), simulation of the effect of small crowdants on mean MT lengths (Figure S3B), as well as single filament dynamics reported in literature^21^. Thus while bulk viscosity increases uniformly with crowdant concentration for both LMW and HMW crowdants (Figure S1A,B), we demonstrate that elongation rates depend on crowdant size. LMW crowdants like glycerol and ethylene glycol decrease elongation rates, while HMW crowdants increase them as seen for PEG 4000 or do not change them as with ficoll 400 (Figure 9A). We understand this in terms of the differential effect on micro-viscosity as predicted by simulations (Figure 3D). On the other hand the rate of *de novo* polymerization shows an increase for both both LMW crowdants such as Glycerol and ethylene glycol, at different rates (Figure 9B) and HMW crowdants such as PEG 4000 and Ficoll 400. The effect of increasing LMW crowdants thus changes, depending on whether polymerization is in presence of seeds (elongation) or their absence (*de novo* polymerization), and it depend on the combined effect of the number of filaments and their lengths.

**Figure 7:**
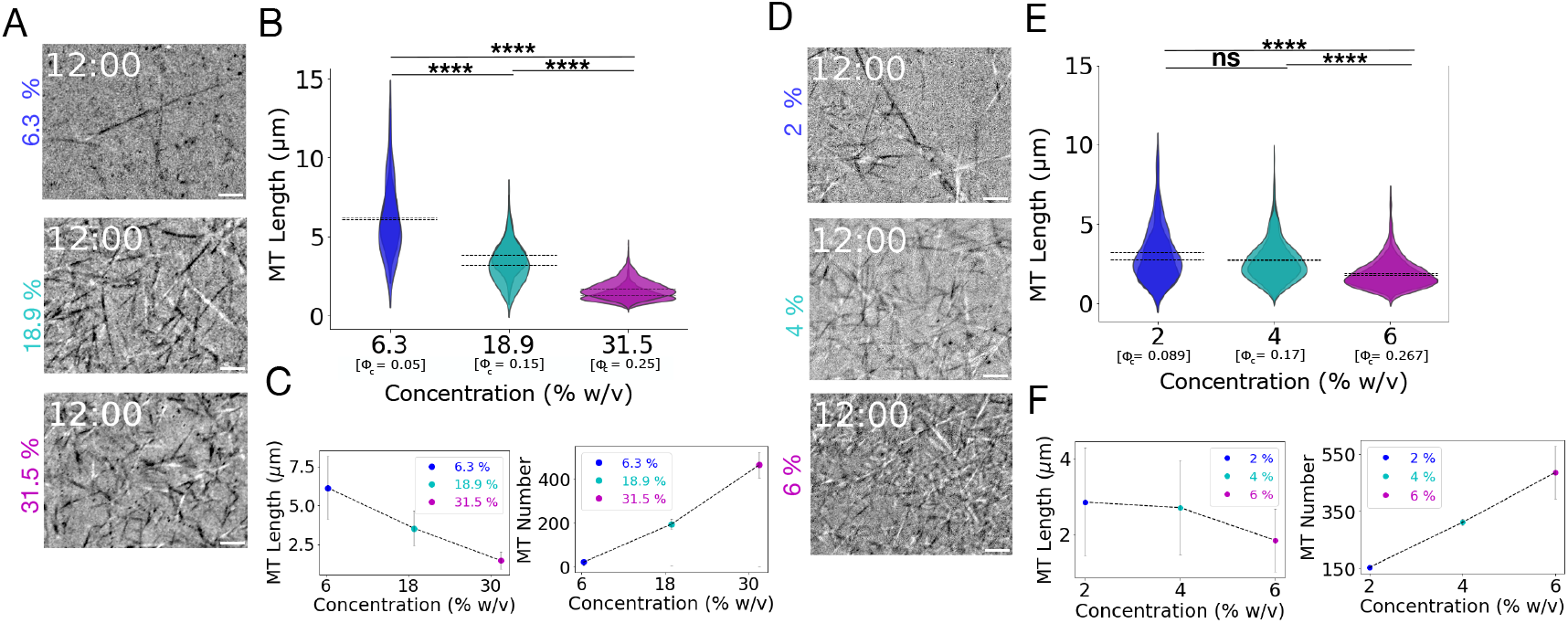
Effect of crowdant concentration and size on MT length distri-bution. (A,D) Representative images from an IRM (label-free microscopy) time-series of 20 *µ*M tubulin with 1 mM GTP polymerization in a coverslip-chamber in presence of (A) 6.3 %, 18.9 % and 31.5 % (w/v) of glycerol or (B) 2 %, 4 % and 6 % (w/v) of PEG 4000 after 12 min. Scale bar: 2 *µ*m. (B,E) The MT length distribution in presence of increasing concentrations of (B) glycerol and (E) PEG 4000. Horizontal lines denote means for individual biological replicates. The differences in mean MT lengths across concentrations were tested for significance using the Mann Whitney U test with **** representing P value *<*0.0001 and ns: not significant. *ϕ_c_*: packing fraction for individual crowdant concentrations. (C,F) Effect of concentrations of (C) glycerol and (F) PEG 4000 on MT Length (left) and MT number (right) respectively. Circles: mean*±* standard deviation, n=6.

**Figure 8:**
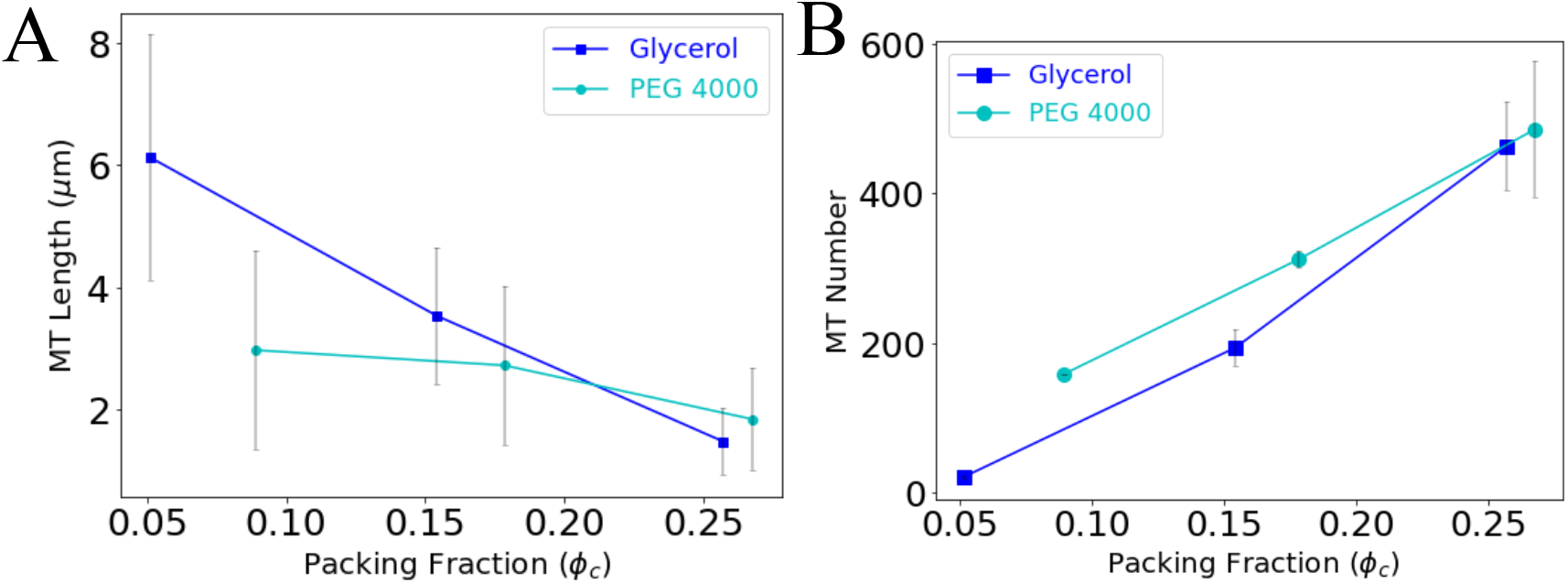
Effect of packing fraction on MT length and number during *de novo* polymerization. The effect of increasing packing fractions (*ϕ_c_*) for two crowdants glycerol (blue box) and PEG 4000 (cyan circles) are quantified from images in terms of (A) MT length and (B) numbers from IRM images in Figure 7. Numbers of MTs analyzed are: (a) Glycerol: n = 41 (6.3 %), n = 387 (18.9 %) and n = 927 (31.5 %), (b) PEG 4000: n = 381 (2 %), n = 622 (18.9 %) and n = 971 (6 %). Symbols represent mean*±*s.d. from two biological replicates.

**Figure 9:**
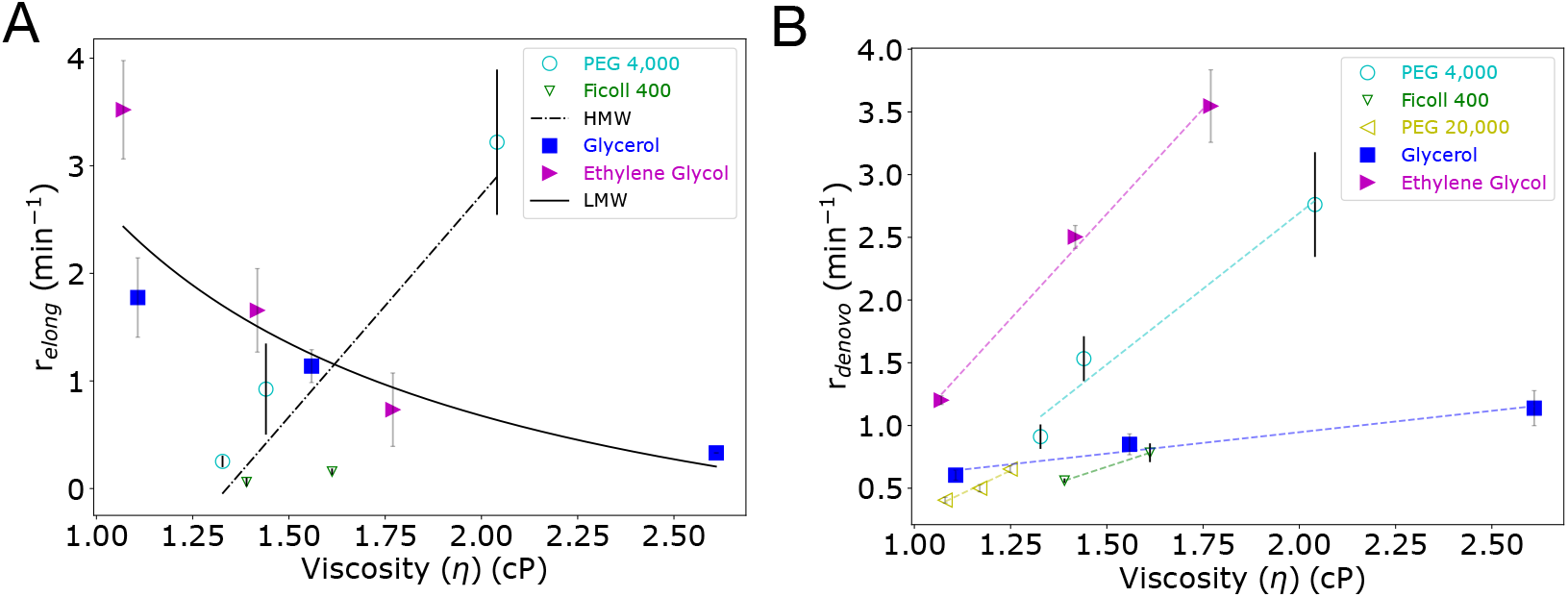
Comparing the effect of viscosity on seed-based and *de novo* polymerization rates *r*. (A) Seed-based polymerization (i.e. elongation) and (B) *de novo* MT polymerization rates *r* from bulk turbidimetric assays were plot-ted against viscosity (cP) for LMW crowdants glycerol and ethylene glycol (closed symbols) and HMW crowdants PEG 4000, Ficoll 400 and PEG 20000 (open sym-bols). The rates of (A) MT elongation, *r_elong_*, and (B) *de novo* polymerization, *r_denovo_*, are obtained from Figure 2 and Figure 5 respectively while viscosity is estimated from microrheology of 1 *µ*m beads in solutions of each of the crowdants (Figure S1). The elongation rate data was fit for large (HMW) crowdants to a saturation function (Equation 6) and for small (LMW) crowdants to *r* = *c*_1_*/η* (Equation 7). The *de novo* polymerization data was fit for each crowdant to a straight line (*r_denovo_* = *c*_2_ *· η* + *c*_3_, where *c*_2_ and *c*_3_ are fit parameters). The R^2^ values were *≥* 0.92. All symbols are mean*±*s.d. from n=6.

## Discussion

Here, we have used a combination of experiment and simulation to test the effect of macromolecular crowdant size and concentration on MT polymerization. We find elongation rates measured by following MT polymerization in presence of seeds are reduced by LMW (small) crowdants while HMW (large) crowdants increase it. These findings at multifilament level are consistent with previous single filament dynamics data and suggest a diffusion limitation of microtubule elongation. We simulate the effect of crowdants on monomer diffusion and MT polymerization using a lattice based kinetic Monte Carlo model and find that (i) monomer diffusion coefficients are reduced more distinctly by small sized as compared to large sized crowdants (Figure 3D), (ii) this is predicted to result from an increase in microviscosity due to LMW but not HMW crowdants and (iii) LWW but not HMW crowdants reduce MT elongation rates (Figure 3E) and mean lengths of polymers (Figure S3B). To test if *de novo* polymerization (in the absence of seeds) is similarly affected by crowdant sizes, we estimated the effect of crowdant concentration and size on critical concentrations, c* in experiment. Crowdants decrease in the critical concentration c* in a concentration dependent manner, independent of their size. We find this is the combined result of LMW crowdant concentration dependent increase in MT number but decrease in MT length, while HMW crow-dants also resulted in an increase in MT number but lengths remained unaffected, based on label free time-lapse microscopy. Our *de novo* multi-filament polymerization experiments demonstrate that the effects of crowdant size on nucleation and elongation appear to act independently-size independent increase in nucleation, but size dependence of MT elongation. The filament microscopy suggests that the increase in polymer fraction seen in bulk kinetics independent of crowdant size is a collective outcome of the effects of crowdants on MT nucleation and elongation.

Microtubule nucleation and elongation are highly regulated in cells by a range of microtubule associated proteins (MAPS) that continue to be discovered (reviewed by Bodakuntla et al.^33^) as well as the ‘tubulin code’ of post translational modifications (reviewed by Janke and Magiera^34^ and Roll-Mecak^35^). At the same time, the physical properties such as macromolecular crowding also affect microtubule polymerization, and are important since the cell is a crowded environment. Previous studies had established that the critical concentration of brain tubulin polymerization could be reduced from the *∼*10 *µ*M range under standard conditions^30^ to 2.5 *µ*M on addition of crowdants like polyethylene glycol 6000 and dextran-T10^36^.

Glycerol was also shown to increase the rate of initial MT assembly and slowing depolymerization^37^. The effect of small crowdants to decrease MT elongation rates of microtubules while large crowdants showing increased elongation has been reported before^20^ and discussed in terms of *in vivo* viscosity increases^21^. However, as we can show the viscosity measured using micron sized spheres increases irrespective of crowdant size (Figure S1). This divergence of effects of bulk viscosity on polymerization depending on crowdant size has been previously been reported for actin elongation rates in presence of LMW while it leads to an increase in elongation rates if the crowdant is HMW^18^. This can be resolved by ignoring bulk viscosity and considering the spatial structure that crowdants generate on the scale of the monomers, with small crowdants resulting in smaller pore sizes as compared to large crowdants^38^. In case of actin this has been demonstrated to increase microviscosity in case of small crowdants thus reducing polymerization rates while large crowdants act through volume exclusion to enhance actin polymerization^19^. In our study we demonstrate using lattice based diffusion simulations that indeed mobility is reduced by LMW but not HMW crowdants, and this is consistent with the diffusion limited nature of elongation. Our results highlight the generality of the results and could help to reconcile potentially contradictory findings.

In a study of the effect of osmolytes and MMC on MT polymerization, they observed a range of 10 to 20% w/v of sucrose (MW 342.30 Da) concentration did alter the *de novo* polymerization kinetics of tubulin^39^. Since the radius of gyration of sucrose is reported to be 0.4 nm^40^, based on our model we expect it to act as a viscogen. In such a case, increasing concentration should lead to increase in polymerization kinetics rate. We believe the reason for the lack of any effect is the range of concentrations used that correspond to packing fractions *ϕ_C_* ranging between 0.05 and 0.09. These are too low to have a significant effect, as per our calculations. Thus, we believe in our work we demonstrate a range of packing fractions that are required to result in some effect on polymerization of tubulin. This could help better understand both *in vitro* and *in vivo* effects observed.

The simulations described here predict a decrease in elongation of MTs with increasing LMW crowdants, consistent with experiments. However, in simulations we do not observe the increase in elongation as a result of increasing HMW crowdant concentration, seen in experiment. In the past explanations for increased rates with crowdants have been attributed to excluded volume interactions (EVE)^14,41^, that serve to increase local concentrations for example during actin polymerization^19^. A possible explanation for the lack of EVE induced rate increases in simulations, that we do observe in experiment, could be due to structure formation of crowdants, that is not explicitly modeled. In future we will seek to further explore models of corrals to represent microdomains ^17^. At the same time, our calculations have allowed for the interpretation of the polymerization and with the addition of multi-step nucleation kinetics^23^ could be used in future to examine more complex *in vivo* scenarios.

Our experiments confirm that MT nucleation appears to be rate limited, while elongation is diffusion limited. This is similar to reports from actin polymerization^18^. Thus MT nucleation showing lowering of critical concentrations in presence of small and large crowdants can be understood to arise from excluded volume effects (EVE) while elongation is reduced when viscosity increases due to small crowdants, similar to the observations with actin^18,19^ and amyloid fibrils^14^. At the same time we demonstrate that both bulk viscosity increases as a function of crowdants irrespective of size, for a comparable range of volume packing fraction. It is the role of LMW crowdants in increasing the microviscosity that leads to decrease in elongation rates, by affecting monomer diffusion. This appears to make the interpretation of the *in vivo* data all the more puzzling since Molines et al. ^21^ have reported a decrease in MT elongation rates in *Schizosaccharomyces pombe* cells, due to osmotic increase in intracellular crowding, suggesting the crowding is primarily influencing microviscosity. It would be useful to test whether mixed crowdants *in vitro* would show a dominance of the small over large crowdants to possibly resolve this question.

We conclude that macromolecular crowdant size and concentration influence both MT nucleation and elongation and that our model demonstrates the effect of crowding on diffusion limited MT elongation, while the effect on *de novo* polymerization can be understood as a collective outcome of the increase in nucleation independent of crowdant size, with a size-dependent change in elongation rates. This would could help better understand the role of such collective physical effects on the growth and regulation of cytoskeletal polymers in cell physiology.

## Material and methods

### Purification and polymerization kinetics of brain tubulin

Tubulin was purified from freshly sacrificed goat brains as discussed by^26^ modified from a temperature based polymerization depolymerization method described by^42^. Purified brain tubulin polymerization kinetics were measured by measuring absorbance at 340 nm using 96-well half-area UV-transmissible flat bottomed plates (Corning, USA) in a micro-titer plate reader (Varioskan Flash, Thermo Scientific, USA) with 20 to 60 s time interval typically over 50 minutes. Two kinds of experiments were performed: (a) MT elongation kinetics and (b) *de novo* polymerization kinetics, both in the presence of a range of macromolecular crowdants with divergent molecular weights: glycerol (MW 92 Da), polyethylene glycol (PEG) 4000 (SRL,India) (MW 3.5 to 4.5 kDa), Ficoll 400 (MW 400 Da) and ethylene glycol (MW 62.07 Da),polyethylene glycol (PEG) 20,000 (SRL,India) (MW 3.5 to 4.5 kDa) and bovine serum albumin (BSA) (MW 66.43 kDa).

#### MT elongation kinetics

Stable MT nucleation seeds were prepared by incubating 30 *µ*M tubulin with 0.8 mM GMPCPP (NU-405L; Jena Bioscience, Germany) in BRB-80 buffer (80 mM PIPES, 1 mM EGTA, and 1 mM MgCl_2_) at 37 *^o^*C for 30 to 40 minutes, and till the absorbance at 340 nm became stable. The assembled MTs were then sheared by ultrasonication (Ultrasonic Cleaner Sonicator, Spire Automation & Innovation India) for 1 minute. The seeds were further sheared by pipetting using 10 *µ*l tips and their presence confirmed in IRM microscopy. 15 *µ*l of seed solution was mixed with 25 *µ*l of tubulin monomer solution (8 *µ*M tubulin in 1 mM GTP, 5 mM MgCl_2_ and BRB-80 along with a crowdant) and absorbance at 340 nm monitored at 37*^o^*C for 30 to 50 mins, at an interval of 60 s.

#### De novo polymerization kinetics

A 30 *µ*l reaction mix was prepared containing 30 *µ*M tubulin, 1 mM GTP, 5 mM MgCl_2_, BRB-80 and crowdants, and incubated at 37 *^o^*C. Blank samples contained everything except tubulin. Absorbance at 340 nm was monitored at an interval of 20 s.

### Data analysis and fitting

The A_340_*_nm_* values were path length corrected to 1 cm and blank (polymerization solution only lacking tubulin) subtracted. Data from individual experiments was averaged over two independent replicates each with three technical replicates and mean absorbance (A*_t_*) with standard deviation calculated and plotted as a function of time. The absorbance kinetics data was fit to a standard four-parameter model used previously to fit for nucleation limited polymerization kinetics^26,39,43,44^ as follows:

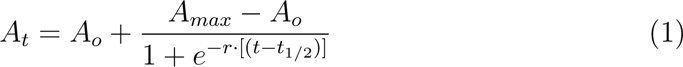

where r is the polymerization rate (min*^−^*^1^), t_1*/*2_ is the time at which absorbance is half-maximal (min), A*_max_* and A_0_ are the maximal saturation and initial absorbance values respectively (arbitrary units).

The normalized steady state absorbance was calculated as the difference between the maximal and minimal absorbance *A_n_* = [*A_max_ − A*_0_], to correct for the initial values that differ between experiments. The steady state absorbance (A*_max_*) was used to calculate the total polymerized tubulin and was used as a parameter for determining the critical concentration. The steady state absorbance and concentration of tubulin was fit to a line as follows:

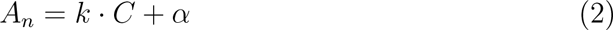

where C is the concentration of tubulin with k and *α* as the slop and y-intercept. The critical concentration (c*) then is the x-intercept of this line, i.e. *c∗* = *−α/k*.

The packing fraction (*ϕ_c_*) in experiment for the crowdants of different molecular weights was determined using the following equation:

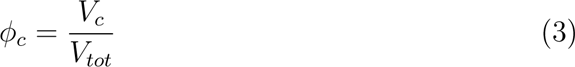

where V*_c_* denotes the volume occupied by crowdant molecules and V*_tot_* denotes total reaction volume, which in this case was 30 *µ*l or 3 x 10^19^ nm^3^. The volume occupied for individual crowdants (V*_c_*) was calculated from the hydrodynamic radius (r*_h_*, in nm) of the crowdants and number of crowdant molecules (N*_c_*) according to the following equation,

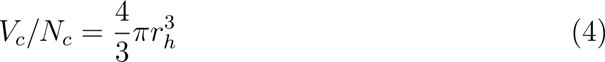

The concentrations of crowdants (C, gm per 100 ml) was used to calculate *N_c_* from the given volume, v= 30 *µ*l, the molecular weight of the crowdant (MW, in gm) and Avogadro’s number (N*_A_*) as follows:

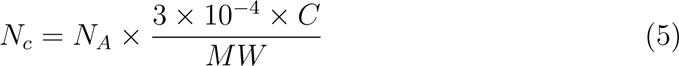

The elongation rates estimated from seed nucleated polymerization kinetics were fit using either of two functions depending on whether the rate increased or decreased: (a) a saturation model based loosely on Michaelis-Menten kinetics for rates that increased with viscosity:

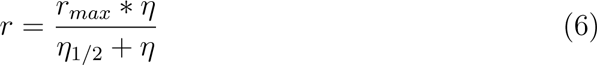

where *r_max_*is the maximal rate and *η*_1*/*2_ is the viscosity where the rate is half-maximal and (b) a function is inversely proportional to viscosity, for decreasing rates:

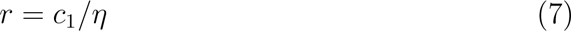

where *c*_1_ was a fitting constant. *De novo* polymerization rates were more heterogeneous due to the combination of the underlying multi-step nucleation and elongation, and were fit to straight lines. All data analysis was done using Python (ver. 3.9.12) with Numpy (ver. 1.21.5) and Scipy (ver. 1.7.3) libraries.

### Bead diffusion microrheology to estimate micrometer scale viscosity in presence of crowdants

Viscosity of different crowdant concentrations were estimated by a microrheological approach using an approach we have previously described elsewhere^45^ that we briefly summarize here. Polystyrene beads of 1 *µ*m diameter (Polysciences inc, USA were diluted in BRB80 buffer along in combination with a range of concentrations of crowdants and mounted in double-backed tape chambers with a circular hole sandwiched between a slide and coverslip. Time-series images were acquired using a 40x (NA 0.65 ELWD) lens using a Nikon TiE inverted microscopy (Nikon Corp., Japan) coupled to a CCD camera Andor Clara2 (Andor Technology, Belfast, UK) every 0.5 s for 1 min. The diffusion coefficient was estimated from fitting the plot of the mean square displacement (MSD) with time using a MATLAB program DICOT^45^ and the viscosity then estimated by substitution in the Stokes-Einstein relation.

### Interference reflection microscopy (IRM) of tubulin filament polymerization

Tubulin polymerization was visualized in microscopic round chamber setups as shown (Figure S5), created using cleaned microscope slide and No 1.5 coverslips with a thickness of 0.17 mm (VWR; Avantor, Radnor, PA). The coverslips were pretreated with 1 N HCl at 50 *^o^*C for 5 hours, followed by rigorous washing with double distilled water to remove any traces of acid, followed by 70 % ethanol wash. 20 *µ*M tubulin in presence of 1 mM GTP, 5 mM MgCl_2_, different crowdant concentration in BRB80 buffer was added into the round chambers created by double backed tape on slides, and were sandwiched by the coverslips.

Samples are imaged using a 100x (NA 0.95 oil) lens on a Nikon Eclipse Ti-E inverted microscope at 37 *^o^*C by a temperature control system (Okolab, Pozzuoli, Italy).Interference reflection microscopy (IRM) images were acquired with a 50/50 beamsplitter (Chroma Technology Corp., Bellows Falls,VT) in the reflected light path based on a previously described method^26,46,47^. Images were acquired every 1.5 mins for 30 mins with an Andor Clara2 CCD camera (Andor Technology, Belfast, UK). Images were background subtracted by taking a median projection of the time-series, applying a gaussian filter and subtracting this ‘background image’ from every frame of the original time series, to enhance contrast. The 32-bit images were then converted to 8-bit for further analysis. All image processing and analysis was performed using the image processing software, FIJI ver.2.3.0/1.53q^48^.

### Lattice-based Kinetic Monte Carlo model of tubulin diffusion and polymerization in presence of crowdants

A rule-based lattice kinetic Monte Carlo model was developed to simulate linear polymer growth by addition of monomers to the ends, with monomers and crowdants diffusing. This approach is comparable to previous studies modeling lattice based diffusion and reaction^17,49,50^. We model four entities: (i) monomers (m), (ii) nucleation points (n), (iii) polymers and (iv) crowdants (c). Each of these entities is modeled to have distinct microscopic events: (i) monomers undergo diffusion, nucleation, polymerization and dissociation from the polymer, (ii) nucleation points can form by aggregation of two or more monomers and depolymerization unless they are pre-seeded, (iii) polymers grow and shrink at fixed rates ends by addition of monomers, with both ends having identical on- and off-rates and (iv) crowdants undergo diffusion. Volume exclusion is implemented for all components of the model, such that a lattice if occupied by an entity, cannot be occupied by another. For simplicity of computation, monomers and crowdants are diffusive but nucleating ‘seeds’ (short polymer segments) and polymers are not. In the following sections we describe the diffusion and polymerization models.

#### Lattice-based diffusion model

Particles can move on a square lattice with 4 possible positions separated by *±δ* in the direction of basis vectors of the lattice. Here, *δ* is spacing between lattice points, while the basis vectors of the square lattice are *{{*1, 0*}, {*0, 1*}}*. Simulations were performed on a finite lattice size of size *L* with periodic boundary conditions. Monomers and crowdants occupy lattice elements of fixed and finite size given by *L_monomer_* and *L_crowdant_* respectively. Mobility is uniform in all directions. Crowdant size is varied to be smaller, equal to and greater than monomer size, in order to test the role of crowdant size on monomer diffusion and polymerization (Table S1). Diffusion is modeled as on-lattice random walks restricted to lattice sites that are integral multiples of the lattice spacing. The diffusive mobility of each molecule (m or c) is based on unit step-length *δ*. Diffusion of monomers and crowdants is modeled by a probabilistic, particle specific, hopping rate *r_D_* = 2 *· d · D_i_/δ*^2^. The dimension of the system *d* = 2 and *D_i_* is the diffusion coefficient of the *i^th^* species (monomer or crowdant of different sizes) based on the Stokes-Einstein relation. The size dependence of either monomer or crowdant diffusion is given by substituting the Stokes-Einstein relation for *D_i_*, where:

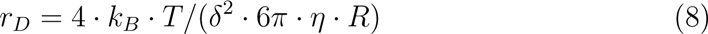

where *R* is the hydrodynamic radius of the molecule, *k_b_* is Boltzmann constant, T is temperature and *η* is the viscosity. The probability of stepping is modeled as a Poisson process with the inter-arrival time modeled based on the expression *E*(*r_D_*) = *−log*(*U* [0, 1))*/r_D_*, using an exponential random number in the domain 0 to 1 as described previously^51^. Random numbers are then generated that satisfy the distribution for a given diffusion rate. The hopping process is then repeated until the cumulative sum of inter-arrival just exceeds the integration time.

#### Macromolecular crowding

Crowding is modeled by volume exclusion, i.e. no two entities in the lattice can occupy the same lattice sites simultaneously. This condition ensures that lattice site occupancy is mutually exclusive, without any overlap. Entities can however share a boundary.

#### Polymer growth

Monomers (*m*) can bind to a pre-existing nucleus (*n*) at either end to form a a polymer (*p*) of length *k* monomer units at a rate given by *k_pol_*, reversibly with a depolymerization rate *k_depolP_* as described the mass action relation:

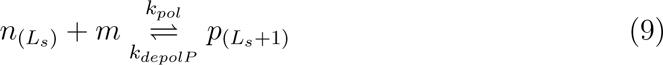

Here, *L_s_* is the length of the seed in monomer units. Monomers can also add to polymers at the rate *k_pol_* and dissociate at *k_depolP_* as follows:

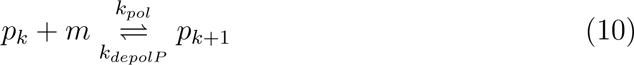

In the model nuclei do not interact directly with polymers. Diffusing monomers (*m*) are tested for interaction after every jump based on proximity to ends of nuclei or polymers. For interaction the contact vicinity of Moore’s neighbourhood of order *L_monomer_* + 1 is chosen where *L_monomer_* is the size of the monomer. If multiple monomers, nuclei or polymer ends satisfy the criterion, one among them is randomly chosen with equal probability.

#### Model parameters and simulations

Model parameters were chosen to capture the relative relations between diffusion of monomers and crowdants and polymerization-depolymerization rates serving as a ‘toy model’ (Table S1). Number of crowdant particles and their sizes were varied, resulting in changes in mobility, in order to test the effect of crowdant size and packing fraction on filament polymerization.

Simulation were performed on Intel’s Xeon Cascade Lake 2.9 GHz processors on a single processor at a time. The system RAM per node used is 192 GB using ParamBramha cluster https://nsmindia.in/node/157#1. Typical simulations run with 2000 monomers and *∼* 40000 crowdants for 200 sec requires 30 minutes.

The code is available on reasonable request to the authors.

## Author contributions

Jashaswi Basu carried out all experiments, data analysis, made the figures and wrote the manuscript. Aman Soni carried out all simulations and made the figures. Chaitanya A. Athale designed and supervised the research, obtained funding and wrote the article.

## Acknowledgments

We are grateful to Shivani Yadav for advice on tubulin purification and polymerization experiments and Thorsten Wohland and Fred Chang for discussions. JB is supported by a PhD studentship from IISER Pune. AS is supported by a fellowship from the Dept. of Biotechnology, Govt. of India (DBT/2021-22/IISER-P/1851) The project was supported by IISER Pune core funds to CAA. The support and the resources provided by PARAM Brahma Facility under the National Supercomputing Mission, Government of India at the Indian Institute of Science Education and Research Pune (IISER Pune) are gratefully acknowledged.

## Abbreviations

• *A*_0_: Initial absorbance at start of kinetics
• *A_max_*: Maximal absorbance at steady state
• *α*: Anomaly exponent of diffusion
• *D_eff_*: Effective diffusion coefficient
• HMW: High molecular weight
• LMW: Low molecular weight
• *L_crowdant_*: Size of a crowdant molecule
• *L_monomer_*: Size of a monomer molecule
• MMC: Macromolecular crowding
• MSD: Mean square displacement
• MT: Microtubule
• NDP: Nucleation dependent polymerization
• PEG: Poly ethylene glycol
• *r*: Rate of polymerization in units absorbance change per minute
• *t*_1_*_/_*_2_: Half maximal time to reach steady state in minutes

## Supplementary Information

1. Supplementary Tables S1 to S3

2. Supplementary Figures S1 to S6

### Supplementary Tables

**Table S1:** Parameters of the monomer diffusion and polymerization model. Parameters of this toy model are chosen to test qualitative trends with values varied relative to each other.

| Symbol | Description | Value | Reference |
| --- | --- | --- | --- |
| *Diffusion model* |  |  |  |
| $\delta$ | Grid spacing | $0.2 \mu\text{m}$ | This study |
| $\eta$ | Viscosity | $10^{-3} \text{ Pa}\cdot\text{s}$ | Viscosity of water |
| $L_{\text{monomer}}$ | Size of a monomer | 3 lattice units | This study |
| $D_{\text{monomer}}$ | Diffusion coefficient of monomers | $0.72 \mu\text{m}^2/\text{s}$ | |
| $L_{\text{crowdant}}$ | Size of a crowdant | 1, 3, 5, 7 and 9 lattice units | This study |
| $D_{\text{crowdant}}$ | Diffusion coefficient of crowdants | 2.17, 0.72, 0.43, 0.31 and $0.24 \mu\text{m}^2/\text{s}$ | " |
| *Polymerization model* |  |  |  |
| $m$ | Initial monomer number | $\sim 2000$ | This study |
| $c$ | Number of crowdant particles | 0 to 40 % | Varied over a range |
| $N_s$ | Number of seeds | 66 | This study |
| $L_s$ | Seed length | 3 monomers | " |
| $k_{\text{nuc}}$ | Nucleation rate | 0 | " |
| $k_{\text{depol}N}$ | Depolymerization rate of nucleus | $0.01 \text{ s}^{-1}$ | " |
| $k_{\text{depol}P}$ | Depolymerization rate of polymers | $0.5 \text{ s}^{-1}$ | " |
| $k_{\text{pol}}$ | Polymerization rate of polymers | $5 \text{ s}^{-1}$ | " |
| *Simulation setup* |  |  |  |
| $\Delta t$ | Time step of simulation | 0.001 s | " |
| T | Total time of simulation | 200 s | " |
| LxL | Lattice size | 420x420 | " |

**Table S2:** Critical concentration for each crowdant concentrations. Crit-ical concentrations (c*) calculated for different concentrations (% w/v) of glycerol (LMW) and PEG 4000 (HMW) as plotted in Figure 6.

| *Crowdant* | *Crowdant concentration (% w/v)* | *Critical concentration, $c^*$ ($\mu M$)* |
| --- | --- | --- |
| Glycerol | 6.3 | 3.85 |
| Glycerol | 18.9 | 2.45 |
| Glycerol | 31.5 | 1.95 |
| PEG 4000 | 2 | 4.88 |
| PEG 4000 | 4 | 3.44 |
| PEG 4000 | 6 | 1.12 |

**Table S3:** Crowdant molecular weights and sizes. Hydrodynamic radius of crowdant molecules is correlated with their molecular weights based on literature.

| *Crowdant* | *Molecular Weight (Da)* | *Hydrodynamic radius ($r_h$)* | *Reference* |
| --- | --- | --- | --- |
| Glycerol | 92 | 0.31 nm | 52 |
| PEG 4000 | $4 \times 10^3$ | 1.92 nm | 53 |
| Ficoll 400 | $4 \times 10^5$ | 7.26 nm | 41 |

### Supplementary Figures

**Figure S1:**
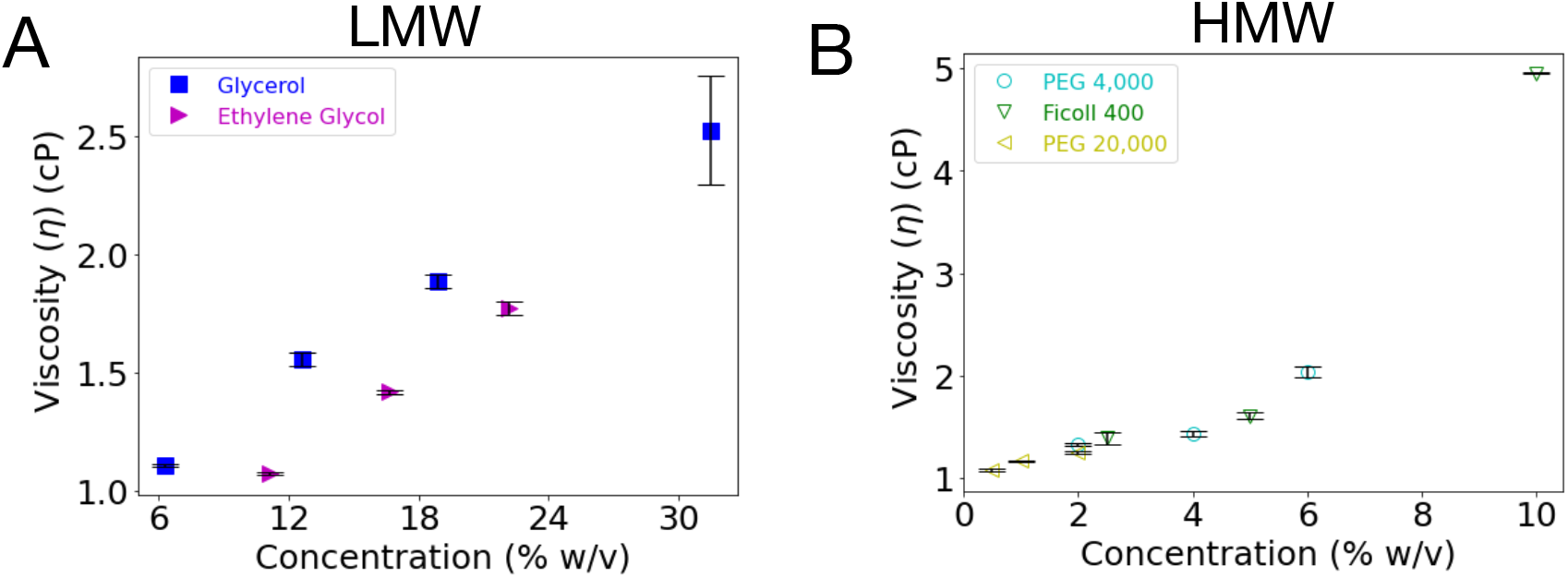
Effect of crowdant size and concentration on viscosity measured by microrheology. Increasing concentrations of (A) glycerol and ethylene glycol (LMW) and (B) PEG 4000, Ficoll 400 and PEG 20000 (HMW) crowdant solutions in BRB-80 buffer were used to measure the diffusive mobility of polystyrene beads of radius 0.5 *µ*m at 37 *^o^*C as described in the Materials and Methods section. The mean square displacement was used to calculate the diffusion coefficient (D), and in turn using Stokes-Einstein’s relation was used to calculate the dynamic viscosity *η* of the solution.

**Figure S2:**
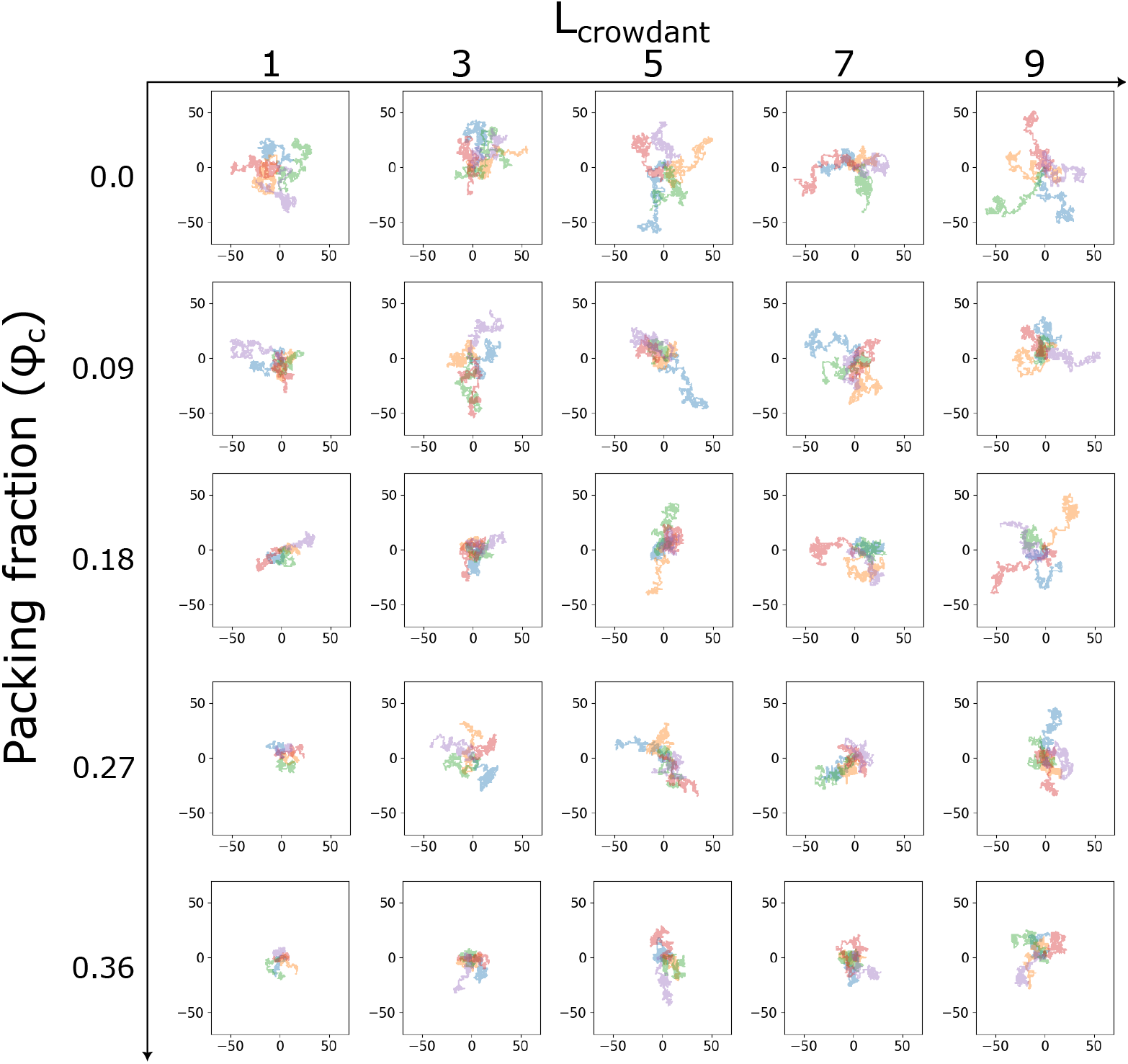
Monomer diffusion simulated in presence of increasing packing fractions of crowdants of different sizes. Monomer trajectories are influenced by varying crowdant sizes and packing fractions within a 424×424 lattice simulation cell. For a visual clarity, ten trajectories over 20 seconds, spanning crowdant sizes *L_crowdant_* = 1 to 9, with increasing packing fractions (*ϕ_c_*). Monomer are uniformly set as *L_monomer_* = 3. The depicted trajectories length shows the effect of different crowdant size and packing into the movement and behavior of individual monomers within the simulation cell. Parameters are listed in Table S1.

**Figure S3:**
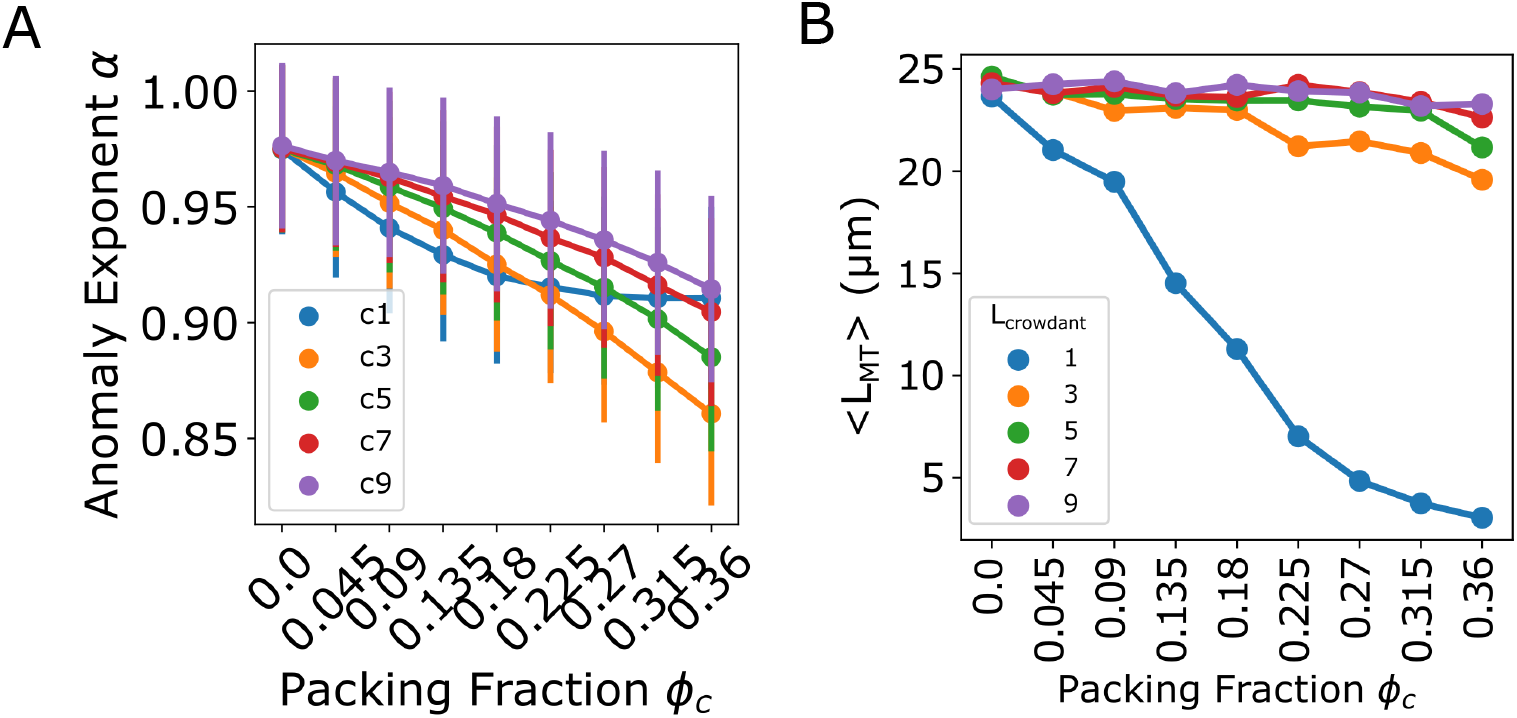
Effect of crowdant size and packing fraction on monomer diffusion and MT length in simulations. (A) The exponent of anomalous diffusion *α* was obtained from fitting the anomalous diffusion model *MSD* = 4*Dt^α^* to the time-averaged MSD of monomers (Figure 3A) simulated as diffusing in presence of crowdants over a range of sizes and packing fractions. (B) The mean filament length from simulations over the same range of crowdant packing fractions and sizes is plotted. Colors: *L_crowdant_* = 1 to 9.

**Figure S4:**
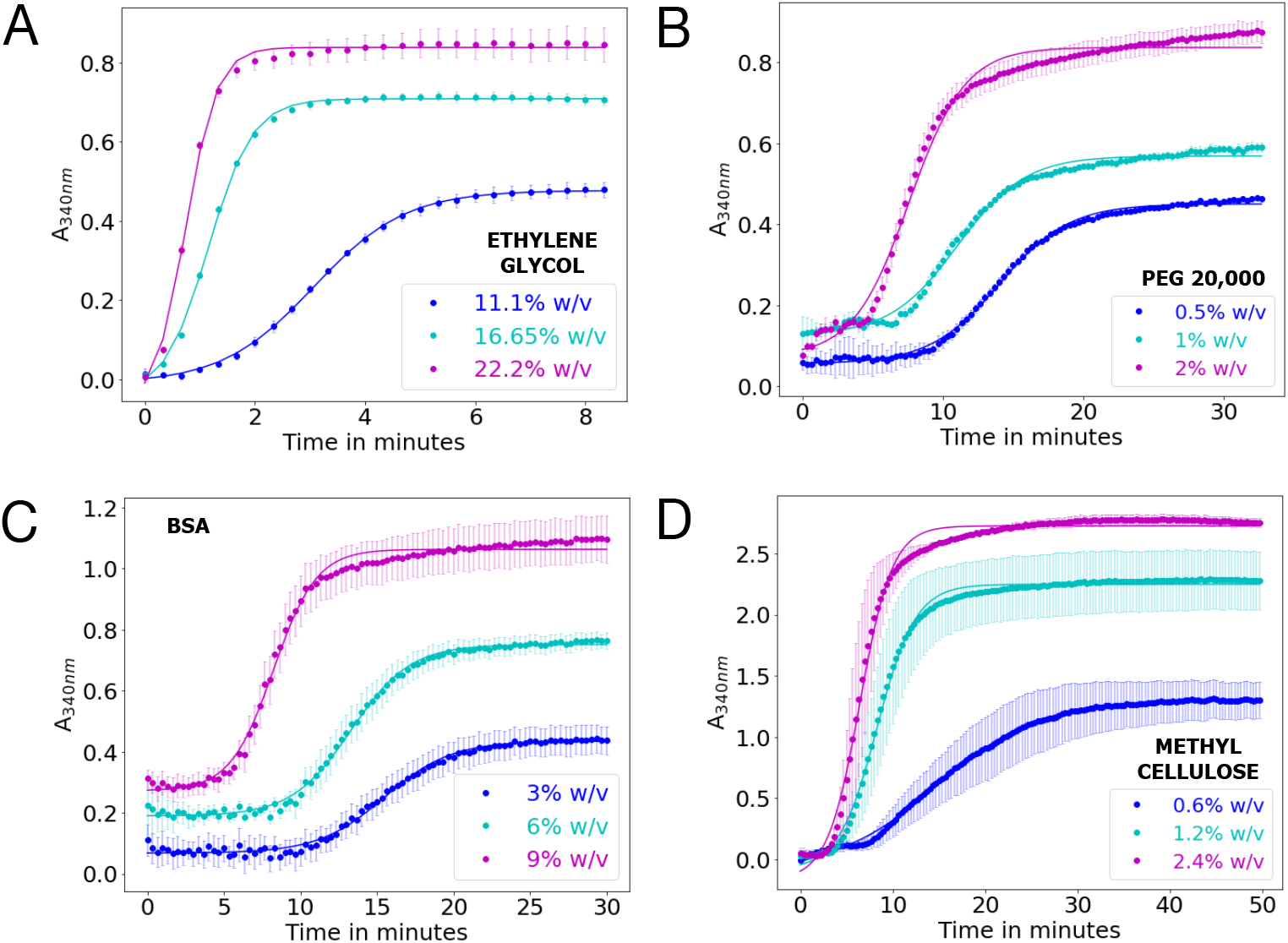
Kinetics of *de novo* tubulin polymerization in presence of increasing concentrations of crowdants. (A) ethylene glycol, (B) PEG20000, (C) BSA and (D) methylcellulose were increased over a range of concentrations (w/v) and the absorbance (A_340_*_nm_*) used to monitor the time dependent kinetics. Colors: crowdant concentration.

**Figure S5:**
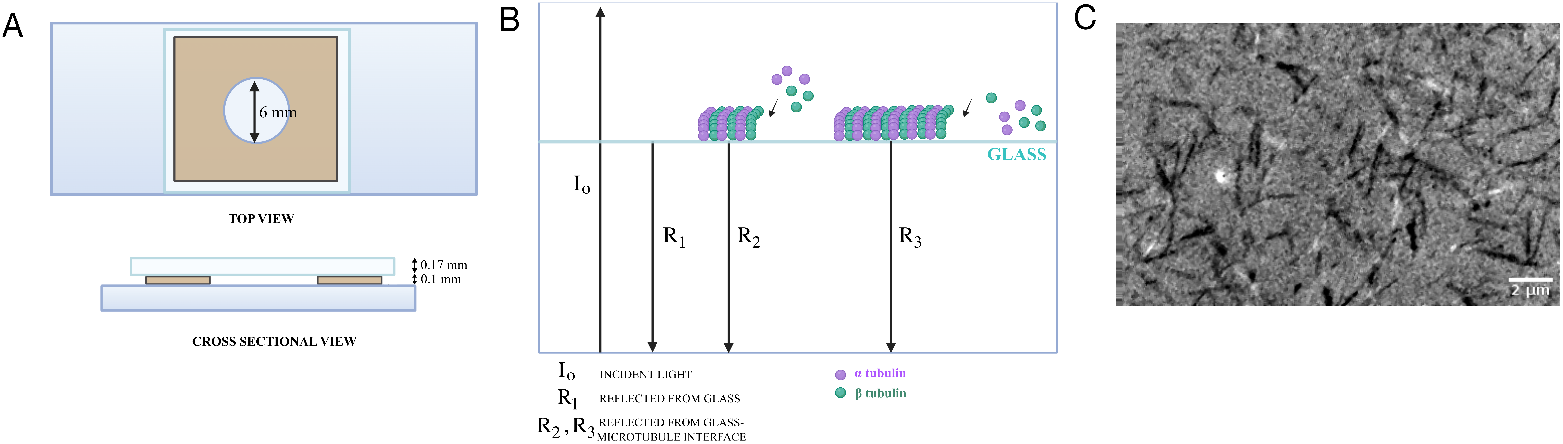
Schematic of microscopy setup to follow tubulin polymerization using IRM. (A) Microscopic slide chamber setup to visualize tubulin polymerization. A round chamber was made with double sided tape (brown) sandwiched by a glass slide and 0.17 mm thick coverslip as seen in top view as well as in cross-section. (B) Schematic of principle by which label-free tubulin polymerization is seen-this figure is adapted from previous reports^46,47,54^. Assembled MTs bind non-specifically to to the glass coverslip. The incident light (I*_O_*) is ei-ther reflected from the glass-water interface (R_1_) or the glass-MT interface (R_2_, R_3_). Image contrast is obtained due the reduced amplitude of R_2_ and R_3_ due to interference. (C) A representative image of MTs polymerized from 20 *µ*M tubulin in presence of 18.9 % w/v glycerol after 3 mins of polymerization is seen after background subtraction, as described in the Methods section. Scale bar: 2 *µ*m.

**Figure S6:**
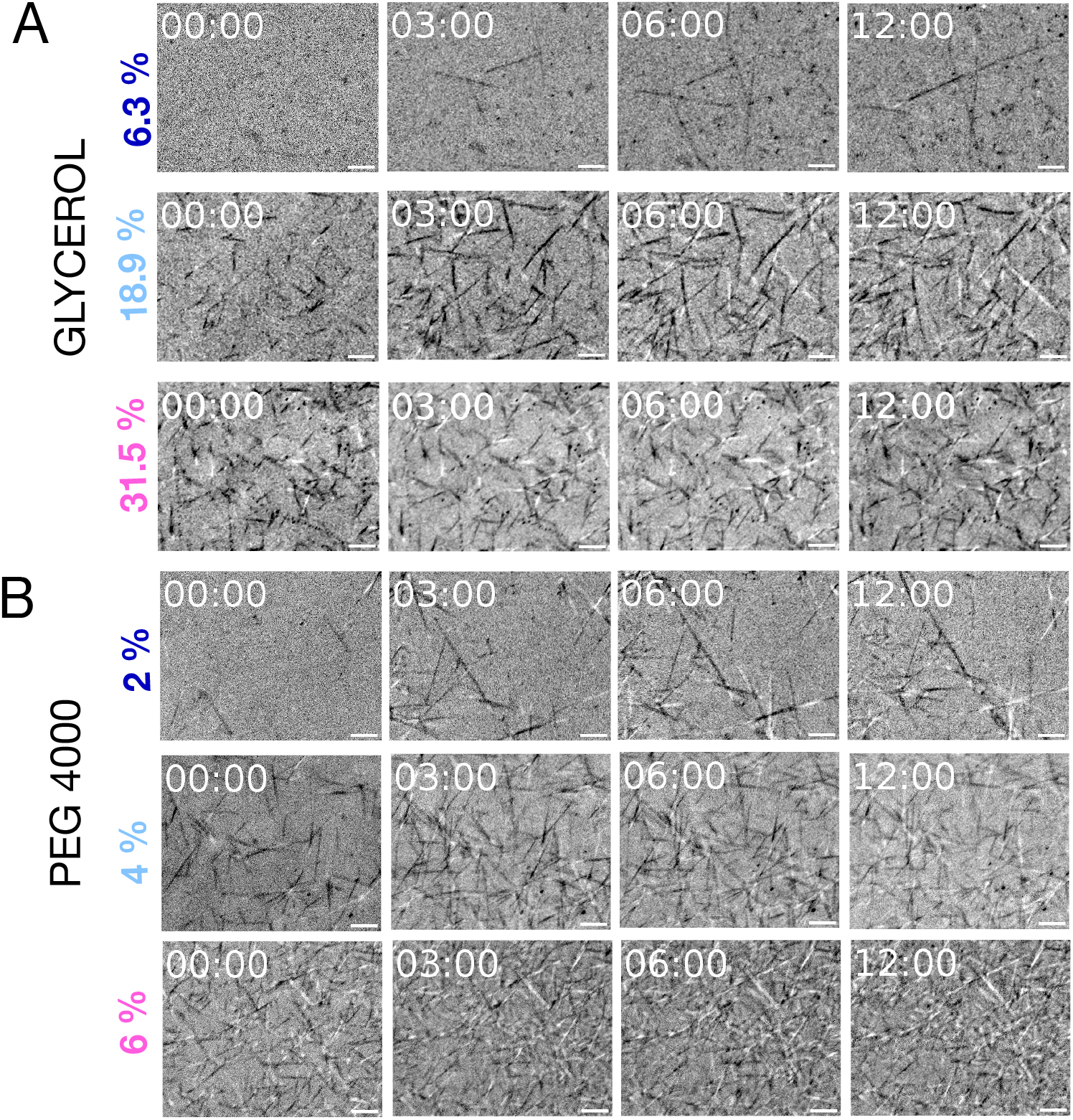
Time-lapse microscopy of MT filament nucleation and growth in presence of crowdants. (A, B) Representative images from label-free microscopy (IRM) time-series of 20 *µ*M tubulin with 1 mM GTP polymerization in a coverslip-chamber in presence of (A) 6.3 %, 18.9 % and 31.5 % (w/v) of glycerol or (B) 2 %, 4 % and 6 % (w/v) of PEG 4000. Time: mm:ss, Scale bar: 2 *µ*m

**Figure S7:**
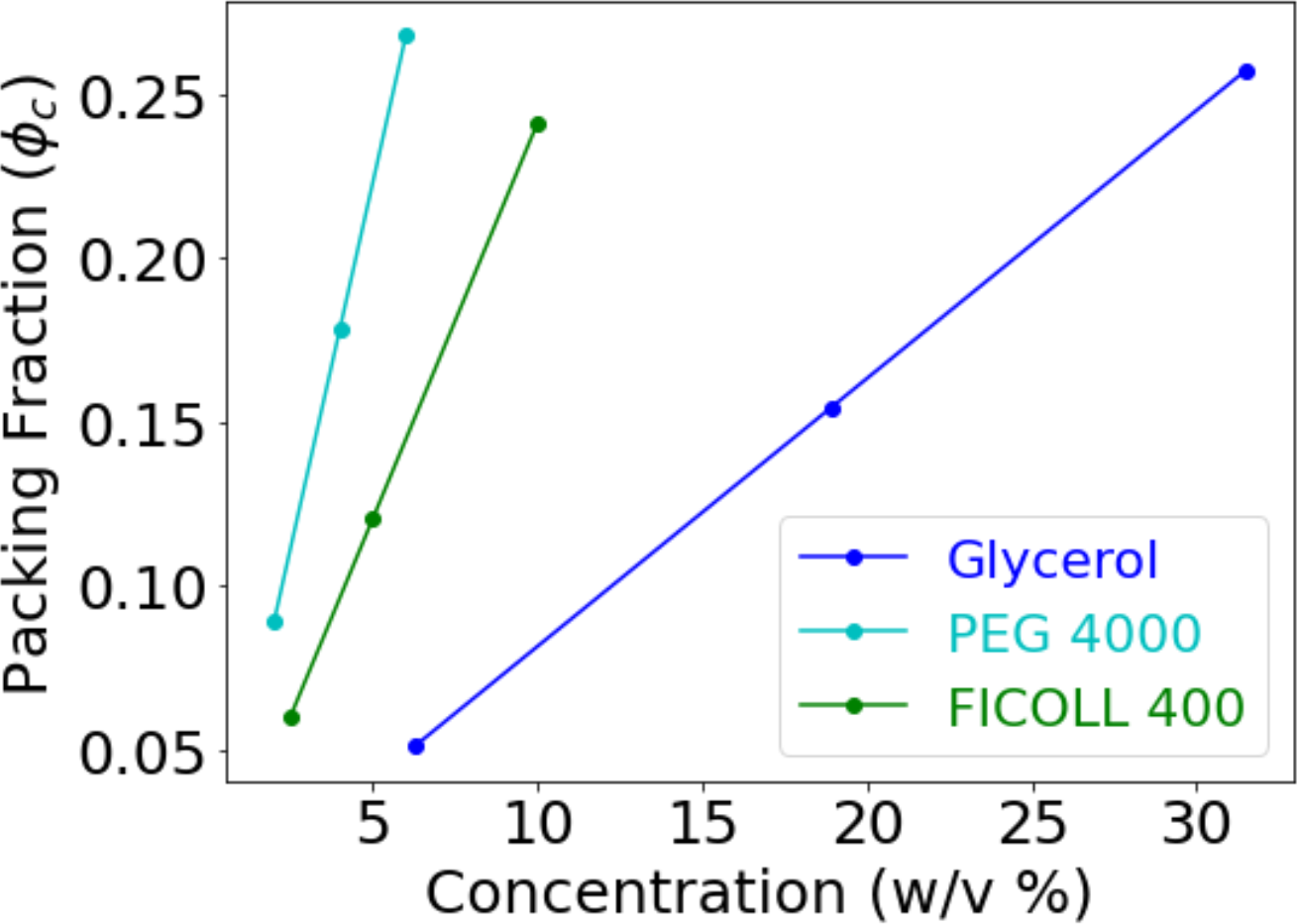
Volume packing fractions (*ϕ_C_*) of crowdants calculated for increasing concentrations. The volume packing fraction *ϕ_C_* was calculated for the crowdants based on the molecular sizes as a function of the % w/v concentrations. Crowdants were: glycerol (blue), PEG 4000 (cyan) and FICOLL 400 (green).

